# Novel symbionts reveal amoebae as significant hosts for environmental chlamydiae

**DOI:** 10.1101/2024.11.01.621453

**Authors:** Fei Liu, Laura Walker, Jason B. Wolf, Christopher R. L. Thompson, Tamara S. Haselkorn, Yijing Shi, Jijuan Ding, Joan E. Strassmann, David C. Queller, Longfei Shu

## Abstract

Chlamydiae represent a diverse group of obligate intracellular bacteria with elusive hosts in environmental settings. This study used one of the largest collections of wild amoebae (*Dictyostelium discoideum* and *D. giganteum,* 106 clones) collected over the past two decades to screen for novel environmental chlamydiae. We found that novel environmental chlamydiae are prevalent in two wild *Dictyostelium* species and assembled 42 novel chlamydiae metagenome-assembled genomes (MAGs). The MAGs represent three chlamydiae species previously only reported using 16S sequencing. Their genomes are divergent enough from other species to warrant placing them in two new genera (tentatively called *Ca. Dictychlamydia sp.* LF1, *Ca. Dictychlamydia sp.* LF2, and *Ca. Feichlamydia sp.* LF3). In addition, these chlamydiae species show strong host specificity with two *Dictyostelium* amoeba hosts, except one amoeba sample. *Ca. Dictychlamydia sp.* LF1 and *Ca. Feichlamydia sp.* LF3 was exclusively observed in *D. discoideum,* while *Ca. Dictychlamydia sp.* LF2 was found only in *D. giganteum*. Phylogenetic and comparative genomic analyses suggest that all three chlamydiae are close to arthropod-associated chlamydiae and likely have some intermediate characteristics between previously reported amoeba-associated and vertebrate-associated chlamydiae. This study significantly broadens our understanding of the chlamydial host range and underscores the role of amoebae as vital hosts for environmental chlamydiae.

## Introduction

Chlamydiae, a group of obligate intracellular bacteria, have been recognized as significant human pathogens for over a century (1). For many years only one clade, composed of many well-known human and animal pathogens, was recognized. However, in recent decades, our understanding of chlamydiae has expanded considerably, revealing a diverse host range and ubiquity in various environments (2–4). Through co-cultivation approaches, researchers have successfully identified chlamydiae in a wide array of eukaryotic hosts, including amoebae, arthropods, and vertebrates (5–7). Additionally, cultivation-independent studies have shown that chlamydiae are prevalent in environmental samples, such as water, soil, and marine sediments, even though their specific hosts often remain unidentified (2–4). Despite their phylogenetic diversity, these chlamydiae are collectively referred to as environmental chlamydiae in order to distinguish them from classical Chlamydiaceae (1). Given that all chlamydiae, including the environmental strains that have been cultured, are strictly intracellular, these findings raise fundamental questions about the unidentified hosts that contribute to the survival, dispersion, and evolution of environmental chlamydiae.

Many efforts have been made to identify the hosts of environmental chlamydiae (3). The most educated hypothesis is that amoeba, ubiquitous unicellular eukaryotes found in both terrestrial and aquatic ecosystems (8, 9), serve as the elusive hosts for many environmental chlamydiae. While knowledge of many protist groups remains limited compared to prokaryotic microbes due to the complex phylogenetic relationships of the protists and cultivation challenges, they exhibit remarkable diversity across environments and many are known to harbor symbionts (8, 10–13). Several protist hosts of chlamydiae are already recognized, primarily in the amoebae, including *Acanthamoeba sp.* (14), *Hartmanella sp.* (*15*), and more recently, the social amoeba *Dictyostelium discoideum* (12, 16). In fact, most chlamydiae are able to survive and replicate in amoeba cells and co-culturing with free-living amoebae has proven an invaluable tool for studying chlamydiae biology (17). It has even been proposed that amoebae are the actual (although unrecognized) hosts of some chlamydiae that are recovered from diverse animal microbiomes (3). Consequently, amoebae, in particular, are plausible hosts for most species of environmental chlamydiae.

However, it is important to note that current evidence provides limited support for this hypothesis. First, most environmental chlamydiae have not been isolated from amoebae. For example, the family *Rhabdochlamydiaceae*, one of the largest and most diverse groups of environmental chlamydiae, has been detected across a wide range of environments, including soil, freshwater, sediment, and marine habitats (5). Surprisingly, the only known hosts of *Rhabdochlamydiaceae* are arthropods (5, 18–20), rather than amoebae. Secondly, many identified amoeba-associated chlamydiae, such as *Parachlamydia* and *Neochlamydia*, differ significantly from environmental chlamydiae. These differences include genome size, as *Parachlamydiaceae* genomes are considerably larger than those of the arthropod symbionts of *Rhabdochlamydiaceae* (5) and they occupy phylogenetic branches distinct from environmental chlamydiae, the bulk of which belong to the *Rhabdochlamydiaceae* family (2). Consequently, there is insufficient evidence to support the idea that amoebae are reservoir for environmental chlamydiae. A noteworthy study by Haselkorn *et al.* (12) found that novel chlamydiae are common in *D. discoideum*, potentially forming a sister group to the *Rhabdochlamydiacea*e. However, 16S sequence analysis proved insufficient for species-level identification, and the study could not yield detailed phylogenetic inferences or any chlamydial genomes for further analysis. (21).

Based on these studies, we aim to address this knowledge gap. We believe the lack of evidence to support amoebae as reservoir for environmental chlamydiae may be due to technical difficulties with working on amoebae. On the one hand, many environmental amoebae cannot be readily cultivated (11), and very few studies have systematically screened for the presence of chlamydial symbionts on large scales. The majority of amoeba-associated chlamydiae species have been identified from *Acanthamoeba* or related species (22). Therefore, the so-called amoeba-associated chlamydiae may be biased and represent only a small portion of the actual amoeba-associated symbionts. In this study, we used 106 wild amoeba clones based on Haselkorn *et al.* (12), that were possible to be reservoir for novel chlamydiae. These amoebae were collected by the Queller-Strassmann group over the past two decades. Two social amoeba *Dictyostelium* species (*Dictyostelium discoideum* and *D. giganteum*), distantly related to *Acanthamoeba*, were used for this study.

## Results

### Novel chlamydiae genomes recovered from two wild *Dictyostelium* amoebae

We conducted an in-depth investigation using next-generation sequencing techniques, resulting in the assembly of 122 high-quality metagenome-assembled genomes (MAGs) (Table S1). Notably, among these genomes, 41 were identified as chlamydiae, which confirmed the presence of chlamydiae in *D. discoideum* (12). Subsequent analyses allowed us to categorize these chlamydiae into three distinct species using a 95% average nucleotide identity (ANI) threshold (Figure S1). To further investigate the taxonomy of these chlamydiae, we compiled genomes from a comprehensive set of 160 species under *Chlamydiales*, sourced from the Genome Taxonomy Database (GTDB) for rigorous comparative analyses (Table S2). A substantial fraction of these chlamydiae specimens were derived from environmental sources, such as soil, marine and freshwater environments, and were not associated with specific host organisms (Figure 1). Comparing to these known chlamydiae genomes, two of the novel chlamydiae genomes assembled here were classified within the *PALSA-1444* group, belonging to the *Rhabdochlamydiaceae* family. These species were identified as the sister clade to the genus *Rhabdochlamydia*, whose hosts were associated with arthropods (Figure S2), and their relative evolutionary divergence ranged from 91.83% to 91.96%, representing a potential novel genus (23). Therefore, we tentatively named them as *Ca. Dictychlamydia sp.* LF1 and *Ca. Dictychlamydia sp.* LF2. The other species represented a previously unidentified genus within the environmental chlamydiae family, referred to as *FEN-1388* (Figure S3). Our analysis revealed that this novel genus displayed a relative evolutionary divergence ranging from approximately 85.50% to 85.53% when compared to reference genomes. Therefore, we tentatively named them as *Ca. Feichlamydia sp.* LF3. Our results confirmed that there were novel chlamydiae within these amoebae samples, and we also assembled their genomes with high quality, that allow us to infer phylogeny and do additional analyses.

**Figure 1.**
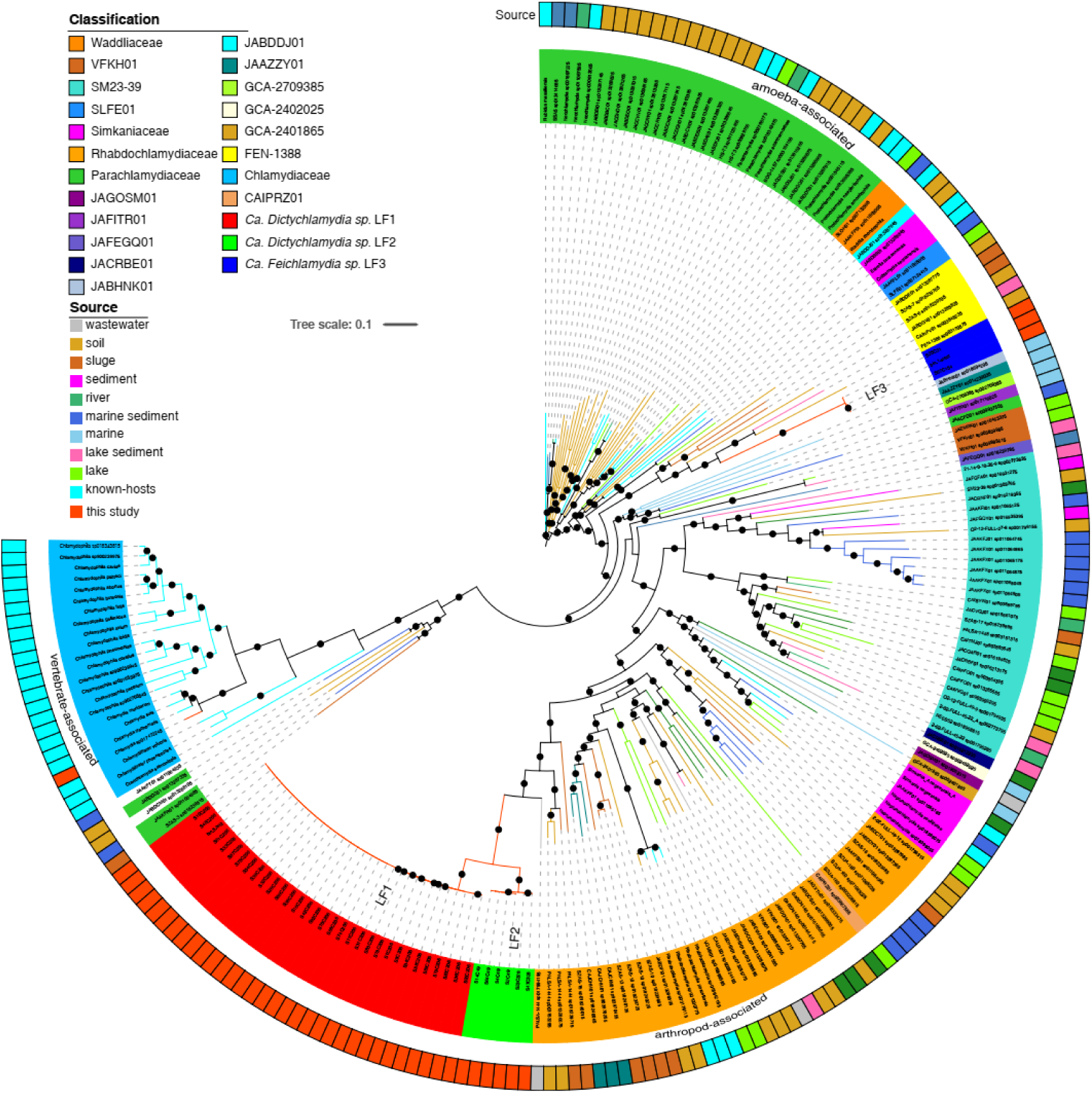
The rooted tree of all 160 chlamydiae from GTDB and 41 novel chlamydiae from this study. The colors on the leaf are the classification of chlamydiae at the family level. The color blocks outside the tree show their source, including the azure blocks representing known-host chlamydiae, and the genomes that obtained from this study mark as red blocks. The points on the tree are the bootstrap (>70). Substitution model: Q.yeast+I+R10.

In the reference genome sets from GTDB database, there were 38 species had clearly defined natural hosts, encompassing 22 chlamydiae species residing in amoebae, 13 in arthropods, and 3 in vertebrates. Notably, one of our previously unclassified species, *Ca. Feichlamydia sp.* LF3, displayed the closest genetic affinity with chlamydiae typically associated with amoebae hosts. In contrast, the other two novel species showed their closest genetic alignment with the *Rhabdochlamydiaceae* family, previously known only to infect arthropods. This finding significantly expands our understanding of the host range of *Rhabdochlamydiaceae*.

### Novel Chlamydiae species show strong host specificity to two *Dictyostelium* amoeba hosts

All wild amoeba clones selected for this analysis were expected to be *Dictyostelium discoideum*, but to our surprise, analysis of 18S rDNA from metagenomic data, indicated that 15 clones are instead, *D. giganteum*. To elucidate the distribution patterns of the novel chlamydiae in the amoebae samples, we conducted an assessment of genomic completeness and sequence coverage across the entire spectrum of wild amoeba clones. The findings unveiled that within our collection of 106 amoeba samples, 42 samples displayed associated chlamydia MAGs with genomic completeness exceeding the 50% threshold, coupled with coverage greater than 1. Remarkably, except for one sample (QS4_A1, *D. giganteum*), our observations were indicative of a predominant association of each amoeba species with a single chlamydiae species, mirroring strong host specificity. *Ca. Dictychlamydia sp.* LF1 and *Ca. Feichlamydia sp.* LF3 was exclusively found within *D. discoideum* (Figure 2A), while *Ca. Dictychlamydia sp.* LF2 was detected in *D. giganteum* (Figure 2B). These findings underscore the distinct host-specific relationships established by these symbionts.

**Figure 2.**
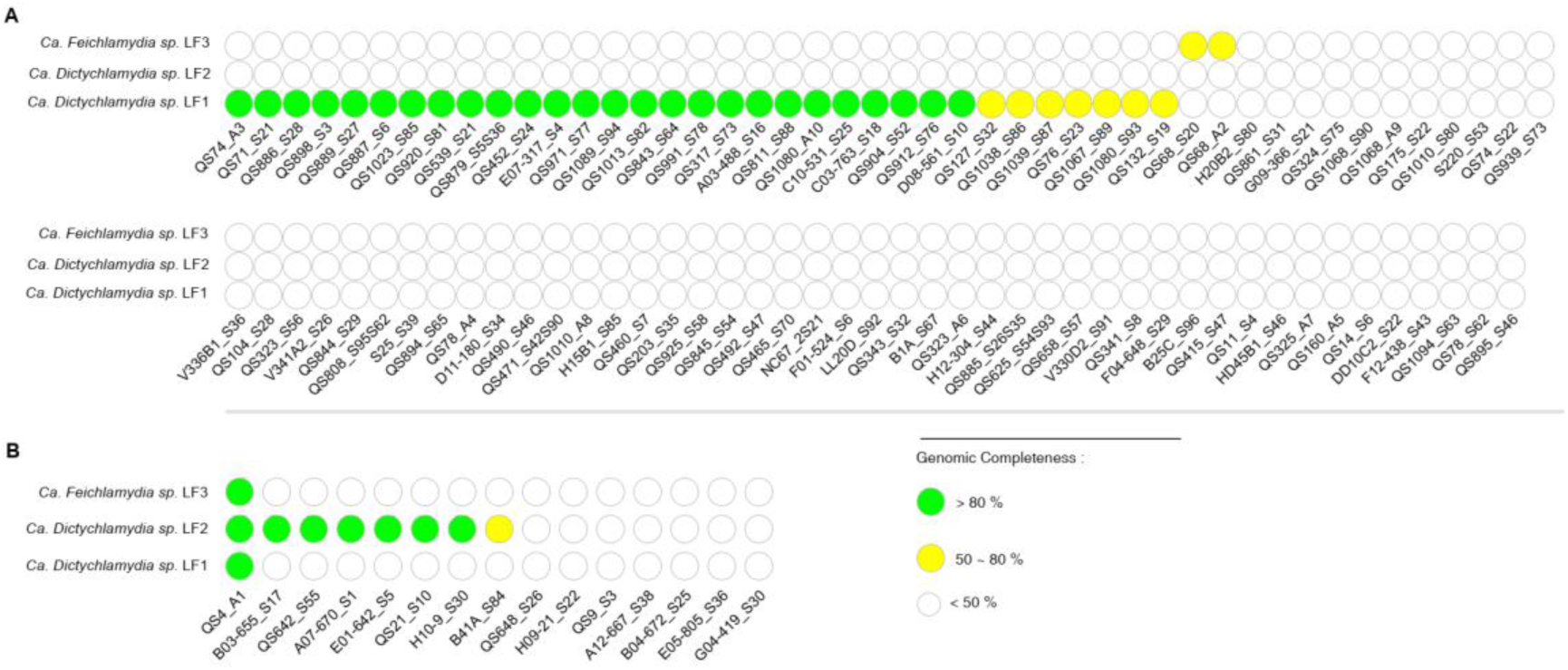
**A:** The distribution of three novel chlamydiae identified in amoebae samples belonging to the *Dictyostelium discoideum*. **B:** The distribution of three novel chlamydiae identified in amoebae samples belonging to the *Dictyostelium giganteum*. Within the two figures: White points: the genomic completeness of chlamydiae less than 50 % within the samples; yellow points: the genomic completeness of chlamydiae ranging from 50% to 80%; green points: the genomic completeness of chlamydiae over 80%.

Interestingly, the distribution of chlamydiae was similar with that in pervious study (12). Our analysis indicated that the *Ca. Dictychlamydia sp.* LF1 and haplotype 1 could be found in the same samples, as well as haplotype 2 and *Ca. Dictychlamydia sp.* LF2. The relationship between *Ca. Feichlamydia sp.* LF3 and haplotype 3 remained less conclusively established, as they were only observed in one same amoeba clone (QS68) (Table S3). Therefore, for further verification, the assembled 16S rDNA, that belong to *Ca. Dictychlamydia sp.* LF1, *Ca. Dictychlamydia sp.* LF2, and *Ca. Feichlamydia sp.* LF3, were used to compared with the sequences of these haplotypes (Figure S5). Surprisingly, the similarity between these 16S rDNA and haplotype 1, 2, and 3 is 99.78%, 99.93% and 100%, respectively. These results provide sufficient evidence to suggest that haplotypes 1, 2, and 3 correspond to the same species as *Ca. Dictychlamydia sp.* LF1, *Ca. Dictychlamydia sp.* LF2, and *Ca. Feichlamydia sp.* LF3, respectively. However, based on the classification, the novel chlamydiae (*Ca. Dictychlamydia sp.* LF1 and *Ca. Dictychlamydia sp.* LF2) were not sister clades of *Rhabdochlamydiaceae*, but belonged to this family.

### Chlamydiae were observed in amoeba spores

Furthermore, we employed transmission electron microscopy (TEM) to validate the presence of chlamydiae within amoeba spores. Four samples, including QS4_A1, QS68_A2, QS74_A3, and QS1080_A10 (with QS4 belonging to *D. giganteum* and the rest to *D. discoideum*), were selected. Chlamydiae endosymbionts were visually characterized as densely wrinkled spheres, measuring approximately 0.5-1 μm in diameter, distributed throughout the cytoplasm (Figure S4). Chlamydiae are known to exhibit multiple morphotypes corresponding to distinct developmental stages (24), notably the infective elementary bodies (EBs) and replicative reticulate bodies (RBs). However, in our observations, a single morphological type prevailed, initially obscuring its precise identity. Clarity emerged when we detected Golgi fragmentation within the amoeba clone (QS68_A2). This observation provided compelling evidence that the chlamydial cells in this particular amoeba sample were in the RB phase. This finding suggests that chlamydiae employ vesicles to exchange materials with their host, as Golgi fragmentation is pivotal in facilitating efficient chlamydial growth (24).

### Phylogenetic and genomic characteristics of novel chlamydiae species

To gain deeper insights into the phylogenetic relationships and genomic attributes of the three newly identified chlamydia species, we selected and analyzed the reference genomes for each species based on the criteria of highest completeness and lowest contamination (Table S4). The genome of *Ca. Dictychlamydia sp.* LF1 (Figure S6) encompassed 1.39 Mbp with a GC content of 41.30% and featured 1145 open reading frames (ORFs). Similarly, *Ca. Dictychlamydia sp.* LF2 exhibited a genome size of 1.68 Mbp, with a GC content of 38.11% and 1261 ORFs, while *Ca. Feichlamydia sp.* LF3 presented a genome of 1.46 Mbp with a GC content of 38.66% and 1265 ORFs (Figure S7). In addition to this, our investigation unveiled 53-83 horizontal gene transfer (HGT) regions within these three species, indicative of ongoing gene exchange events with other bacteria (Figure S8).

Next, we conducted a comparative analysis between the novel chlamydiae and known-host chlamydiae, encompassing 38 reference genomes. Pairwise analysis of whole-genome Average Nucleotide Identity (ANI) revealed considerable variation among chlamydiae, with ANI values ranging from less than 70% to as high as 99.98% (Figure S9). Additionally, we observed noteworthy distinctions in the genome size, Open Reading Frame (ORF) count, and GC content among chlamydiae from different host backgrounds (Figure 3, Table S5). It’s noteworthy that although a correlation between genome size and GC content is commonly observed in many host-associated bacteria, our analysis revealed no such correlation within chlamydiae (Figure 3A). We noted that the GC content and genome size were host-dependent (Figure 3B and 3D), but no such pattern has been previously found in chlamydiae (5). Remarkably, the GC content of the three novel chlamydia species in our study exhibited similarities to chlamydiae known to infect other amoebae hosts (Figure 3A and 3B), whereas their genome sizes were noticeably smaller and more like chlamydiae that infect animals (Figure 3C). From amoebae-associated chlamydiae to arthropod-associated chlamydiae to vertebrate-associated chlamydiae, the genome size and ORF count were consistently decreased. As for the three novel chlamydiae, the results show that they were aligning more closely with arthropod and vertebrate-associated chlamydiae than these amoeba-associated (Figure 3C and 3D), underscoring the impact of host specificity on genomic attributes.

**Figure 3.**
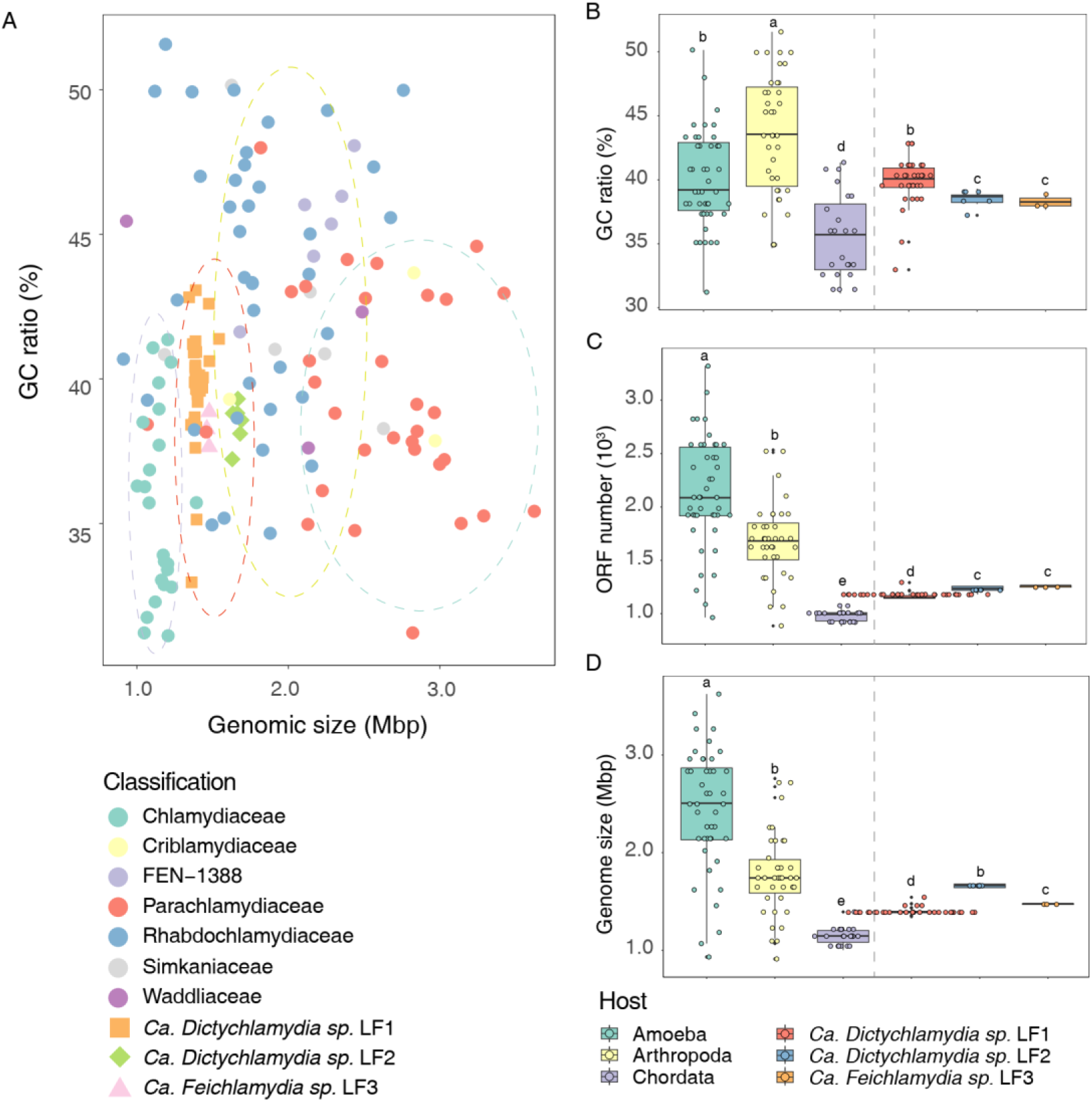
The genomic features of novel chlamydiae. **A:** Correlation of genome size and GC content. The colored circles represent classification at the family level. Other shapes and colors show the 41 novel chlamydiae genomes from this study. **B:** The GC content across hosts. **C:** The ORFs number across hosts. **D:** The Genome size across hosts. The colors represent their host. The letter marks their distinctiveness (ANOVA, *p* < 0.05).

### Functional insights and pseudogene evolution in novel chlamydiae species

Previous research has uncovered remarkable differences in the composition of the outer membrane among various chlamydiae species (25). In our study, we utilized the Transporter Classification Database (TCDB) and the UniRef90 database to annotate genes responsible for encoding outer membrane composition proteins (Figure 4, Table S6). Type III Secretion Systems (T3SS) and Type IV Secretion Systems (T4SS) were found in all chlamydiae species with known hosts. These secretion systems are essential for vectorial secretion and the translocation of anti-host effector proteins (26, 27). Additionally, we observed the prevalence of the *Npt* family, responsible for ATP:ADP exchange, similar to the substrate preferences shown in *C. trochomatis* (28). This suggests that these chlamydiae species rely on their host organisms for ATP and NAD^+^, as they cannot synthesize these essential molecules independently (29). We found that the major outer membrane protein (MOMP) family and NTP (ribonucleoside triphosphates) antiporters were prevalent in chlamydiae with amoeba and arthropod hosts but largely absent in those with vertebrate hosts. In contrast, the polymorphic outer membrane protein (Pmp) family and putative outer membrane porin (OMP) family were more commonly found in chlamydiae species associated with vertebrates. These observations emphasize the host-dependent nature of chlamydiae’s reliance on membrane proteins for ATP and ribonucleoside triphosphates, with distinct transporters used across different host environments. Interestingly, vertebrate-associated chlamydiae do not form the closest grouping; instead, those associated with arthropods are more closely linked with those associated with amoebae.

**Figure 4.**
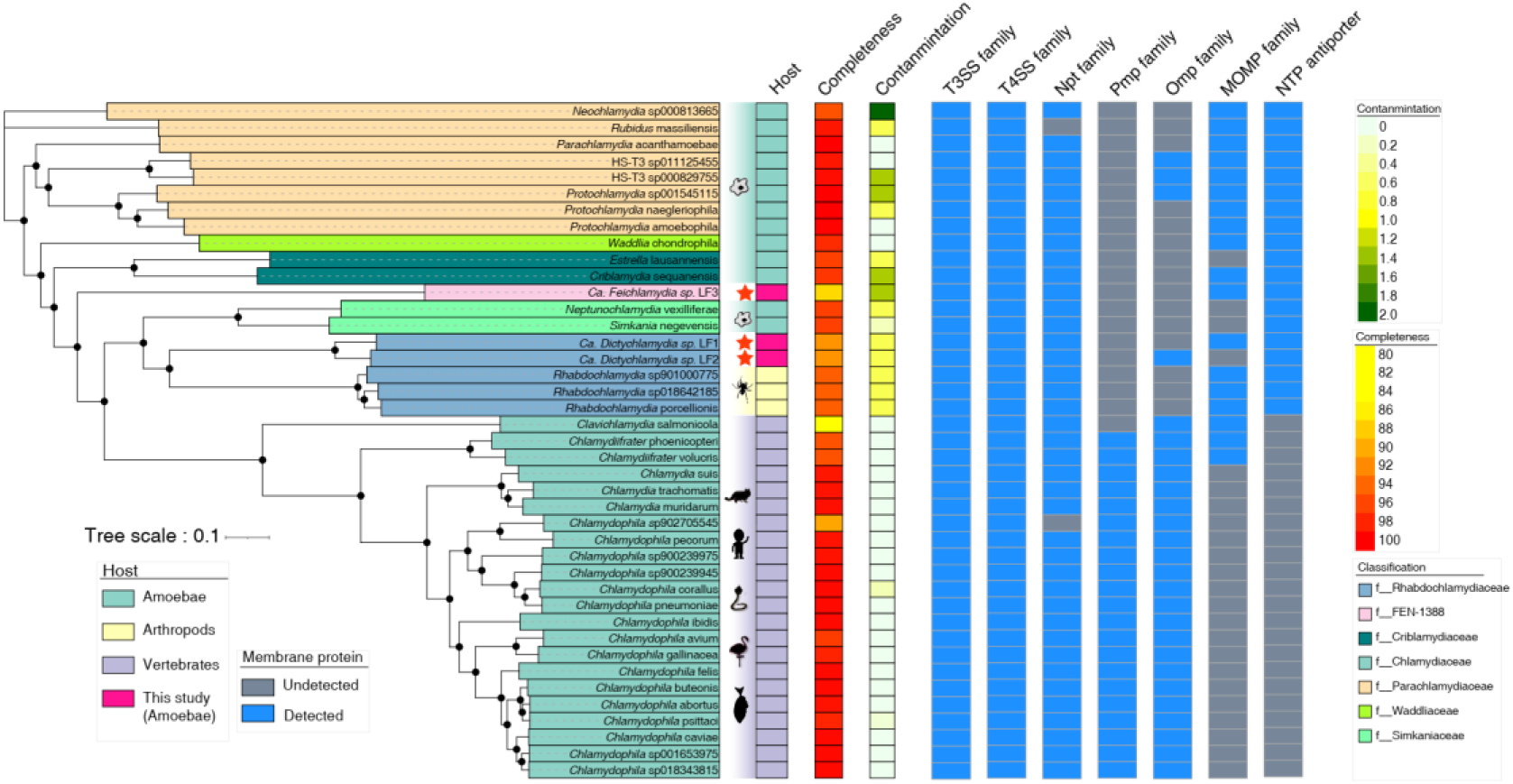
The phylogenetic tree of chlamydiae with known hosts. The colors on the leaf classify chlamydiae at the family level. The black points on the tree are the bootstrap values (>70). Substitution model: Blosum62+F+R4. The blocks represent their hosts, completeness, contamination, and membrane proteins, including type III Secretion Systems (T3SS), Type IV Secretion Systems (T4SS), major outer membrane protein (MOMP) family, NTP (all four ribonucleoside triphosphates) antiporter, polymorphic outer membrane protein (Pmp) family and putative outer membrane porin (OMP) family.

For a comprehensive examination of the functions of the novel chlamydiae, we employed the KEGG database to annotate the open reading frames (ORFs) in these genomes (Figure 5, Table S7). Our results indicated a close connection between chlamydiae’s functional diversity and their host-association. Specifically, the transition from amoeba-associated chlamydiae to those vertebrate and arthropod-associated appear to decreased genome sizes, which might indicate some selectivity. Among functional categories, such as DNA replication proteins and mismatch repair, we observed no changes across different hosts (Figure S10A, ANOVA, *p* > 0.05), totaling 59 genes. In contrast, most functional categories (totaling 688 genes) appear to be decreased trends in arthropod or vertebrate-associated chlamydiae compared to those amoeba-associated, such as the glycine, serine and threonine metabolism related categories significantly reduced functions in arthropod and vertebrate-associated chlamydiae (Figure S10B, ANOVA, *p* < 0.05), while parts of them only decreased in vertebrate-associated chlamydiae, such as ABC transporters related categories (Figure S10B, ANOVA, *p* < 0.05), totaling 214 genes. We also found that functional categories like transfer RNA biogenesis (Figure S10C, ANOVA, *p* < 0.05) were likely to be decreased from amoeba-associated chalmydiae to arthropod-associated chlamydiae, to vertebrate-associated chlamydiae. Other functional categories (totaling 77 genes) exhibited increasing trends, such as riboflavin metabolism (Figure S10D, ANOVA, *p* < 0.05). Importantly, the functional genes of the three novel chlamydiae were more closely aligned with arthropod-hosting chlamydiae, underscoring their possible evolutionary intermediate state (Figure 4).

**Figure 5.**
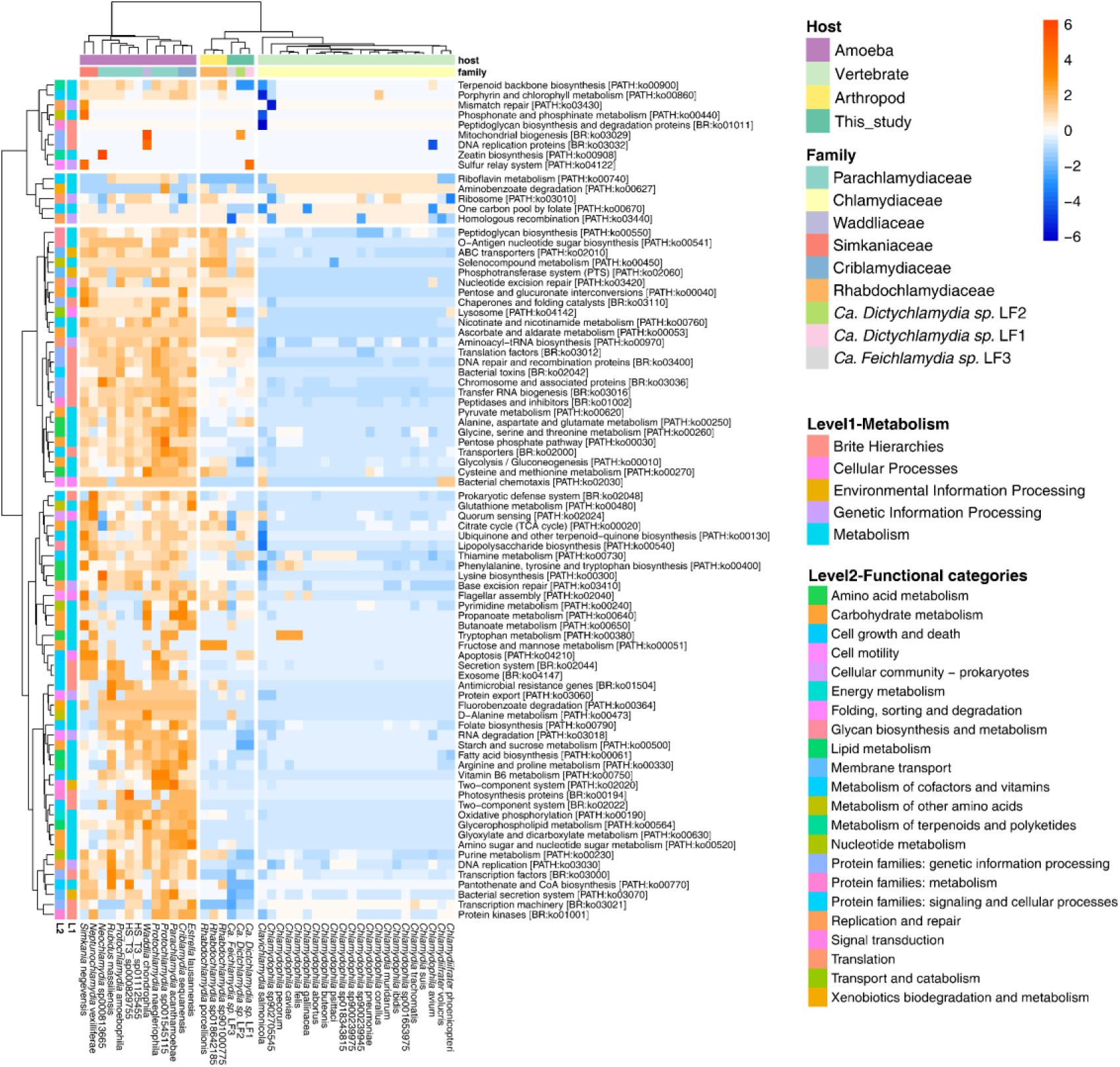
The functions of novel chlamydiae genomes. The y-axis represents functional pathways, and the color blocks indicate different functional classification levels. The x-axis represents species; the color blocks indicate their family-level classification and host information. The heatmap color represents the abundance of functional genes and is normalized by rows. The x-axis and y-axis are clustered by the Euclidean method.

Furthermore, we estimated the potential reasons for the genome reduction observed in the novel Chlamydiae genomes. Similar to previous findings regarding chlamydiae insertion sequences (ISs) (5), our study revealed varying numbers of ISs in the genomes of these three chlamydiae species. Specifically, *Ca. Dictychlamydia sp.* LF1, *Ca. Dictychlamydia sp.* LF2, and *Ca. Feichlamydia sp.* LF3 contained 338, 201, and 139 ISs, respectively. ISs play a critical role in genome adaptation to the host and intracellular environment, often linked with genome size reduction. Parts of genes harboring or near ISs tend to lose functionality, which may lead to their removal during recombination, contributing to genome size reduction (30). Another indicator of genome size reduction is the increased occurrence of pseudogenization, which signifies the degradation of genes within the genome (31). We employed pseudofinder (29) to identify pseudogenes and observed a consistent decrease in pseudogene numbers from chlamydiae hosted by amoebae to those hosted by arthropods and finally to those hosted by vertebrates (Figure 6). Among these pseudogenes, 42.26% were associated with genetic information processing, signaling, and cellular processes (Table S8). Additionally, pseudogenes related to translation and carbohydrate metabolism showed decreasing trends (Figure S11, ANOVA, *p* < 0.05), with their numbers decreasing from amoeba-associated chlamydiae to arthropod-associated chlamydiae and finally to vertebrate-associated chlamydiae. These findings imply that relaxed selection has contributed to deletion mutations and gene degradation, underscoring the correlation between host adaptation and genome size reduction.

**Figure 6.**
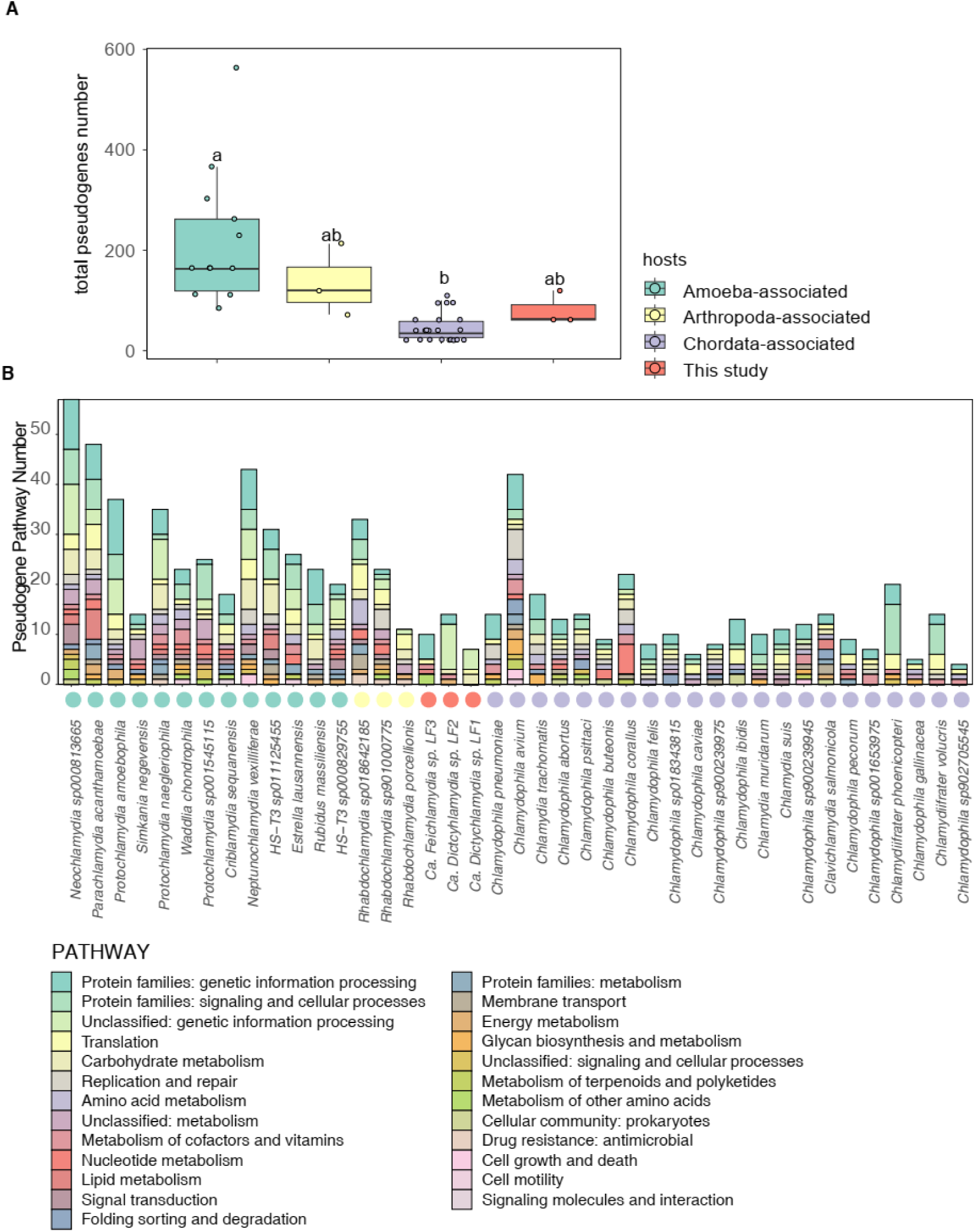
The pseudogenes across chlamydiae of different hosts**. A:** The boxplot shows the difference of total pseudogenes number between amoeba, arthropoda, chordate-associated chlamydia and chlamydiae obtained in this study. **B:** Stacked bar plot shows the annotated pathway of these pseudogenes across their genomes based on KEGG database.

### Metabolic potential of the novel chlamydiae species

Building upon the insights gained from our earlier findings, we reconstructed the metabolic models of the three novel chlamydiae species (Figure 7). Notably, *Ca. Dictychlamydia sp.* LF1 and *Ca. Dictychlamydia sp.* LF2 shared core metabolic functions that were highly similar. Across all three chlamydiae species, we observed an incomplete carbon metabolism and a notable inability to complete the tricarboxylic acid (TCA) cycle independently (Figure 7). These species exhibited complete pathways for fatty acid biosynthesis, although they lacked the capability to utilize fatty acids. We surmise that this pathway may persist as a means to benefit their host organisms. Importantly, these chlamydiae acquired essential resources from their amoeba hosts, including ATP, amino acids, carbohydrates, and various compounds. Intriguingly, our analysis of functional genes revealed distinctions in the ability of the three chlamydiae species to synthesize amino acids. Moreover, we inferred the presence of membrane proteins based on data from the Transporter Classification Database (TCDB) and the UniRef90 database. These proteins play a crucial role in the exchange of signals and nutrients with their host environments. While the precise functions of these membrane proteins remain challenging to ascertain, they are evidently essential for the chlamydiae’s interactions with their amoeba hosts.

**Figure 7.**
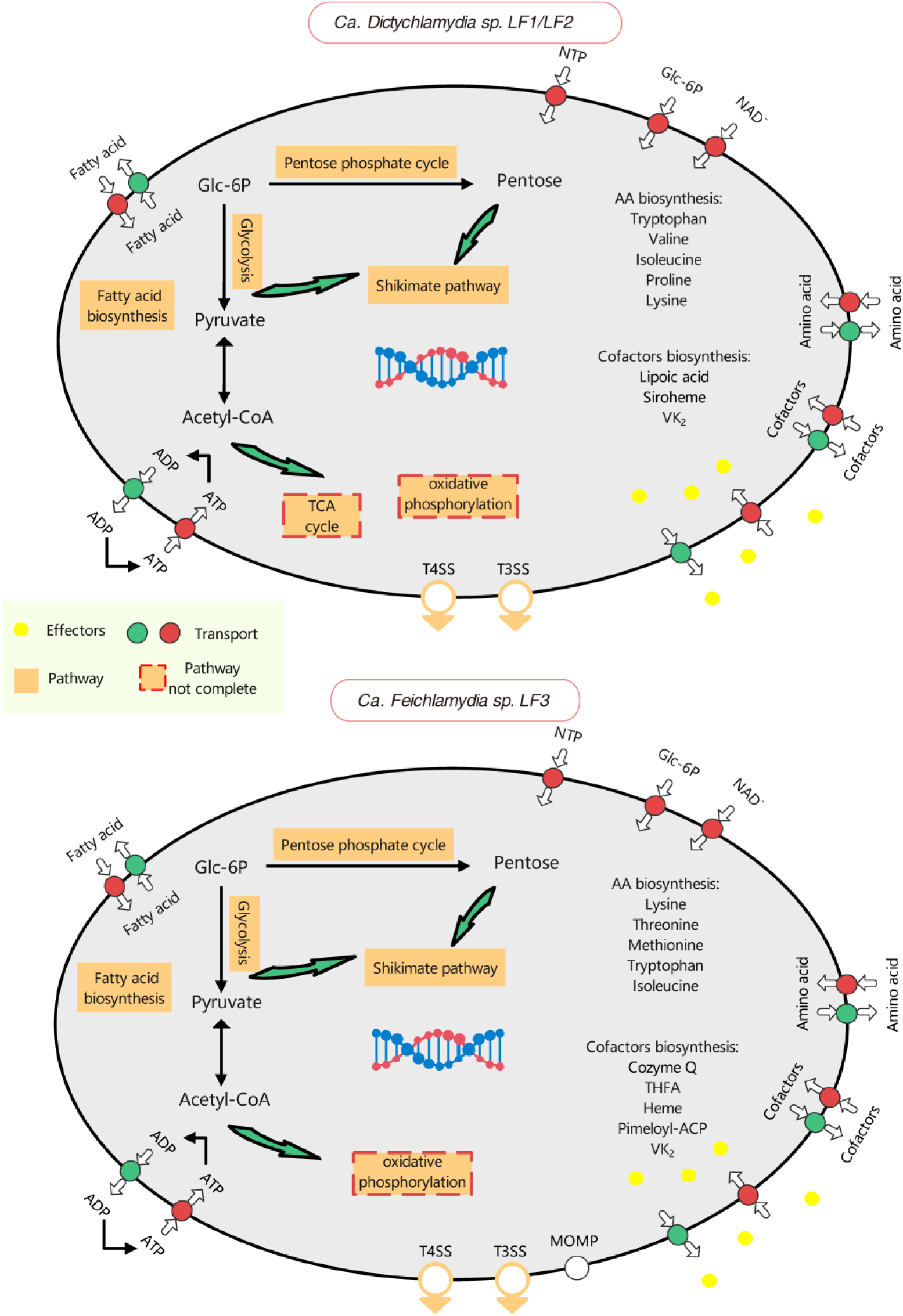
The metabolic models of three novel chlamydia species. The *Ca. Dictychlamydia sp.* LF1 is nearly the same as *Ca. Dictychlamydia sp.* LF2 in the core metabolic pathway we used. Thus, they are shown in the same picture (top figure).

## Discussion

Several species of amoeboid organisms are known to host chlamydiae but much debate remains in the field of chlamydiae biology regarding whether amoebae and other unicellular protists serve as the primary hidden hosts for the diverse environmental chlamydiae (3, 5). A notable study by Haselkorn *et al.* identified endosymbionts associated with wild isolates of *Dictyostelium* amoebae, revealing that 27% of the samples in that study contained 16S haplotypes resembling chlamydiae (12). Based on this study, we selected 106 amoebae samples, that were identified as harbor of novel chlamydiae by Haselkorn *et al*, to investigate the genomes of chlamydiae. Indeed, obtaining a complete genome is much more difficult than amplifying the 16S rDNA, we did not observe the presence of a chlamydia genomes with sufficient confidence in every sample, but a consistent finding between our study and that of Haselkorn *et al*. is that approximately 30% of amoeba samples are infected with chlamydiae. Furthermore, our analysis revealed that novel chlamydiae species show strong host specificity with two *Dictyostelium* amoeba hosts. It is thought that they are related to membrane protein within chlamydiae and their hosts, which have been reported in the Chlamydia (32), and they were called “niche-specific” (33). This helps the evolution of the chlamydia, or even co-evolution with their hosts.

Our results reinforce the notion that chlamydial endosymbionts are widespread in wild amoeba isolates. It is reported that most of known-host chlamydiae can thrive within amoebae except for *Rhabdochlamydiaceae*. However, the *Rhabdochlamydiaceae*, known to exclusively infect arthropods (14, 22), represent a significant portion of the environmental chlamydiae population (5). This leaves a huge gap in hosts of chlamydiae. This study shows that two novel chlamydiae species (belonging to *Rhabdochlamydiaceae*) are prevalent in two *Dictyostelium* amoeba hosts. Therefore, our results show that all major clades of chlamydiae can thrive within amoebae. This study significantly expands our knowledge of chlamydiae’s host range and suggests amoebae as primary unknown hosts for environmental chlamydiae.

This research serves as significant evidence that amoebae play a pivotal role in the evolution of chlamydiae. Functional genes and membrane proteins in amoeba-associated chlamydiae differ notably from those in vertebrate-associated chlamydiae, with arthropod-associated chlamydiae falling in between. Importantly, the genomic characteristics of the three novel chlamydiae species align more closely with those of arthropod-associated chlamydiae. This observation suggests that the novel chlamydiae genomes may represent an intermediate evolutionary state. To further substantiate this, we conducted a phylogenetic analysis using 120 conserved proteins, employing four *Cyanobacteria* as an outgroup (34) to trace the ancestral chlamydiae (Figure S12). Considering that the novel chlamydiae were the clade of arthropod-associated ones, our results propose an evolutionary progression wherein chlamydiae is initially associated with amoebae before transitioning to arthropods and vertebrates.

The gradual reduction in genome sizes from amoeba-associated to vertebrate-associated species and the accumulation of pseudogenes indicate an evolutionary trend towards genome reduction and loss of gene functionality. The genome size of symbiotic bacteria tends to decrease during the evolutionary process (35). Genome size reduction could result from DNA loss and decreased selection to maintain gene functionality (36). We found that the genome sizes of chlamydiae reduced from amoeba-associated to vertebrate, and the others, including the novel chlamydiae, were intermediate (Figure S13). Our results on pseudogenes showed that 42.26% of pseudogenes were related to genetic information processing, signaling, and cellular processes that may not be useful in endosymbionts. Interestingly, some genes gained in the transition to vertebrate-associated hosts, such as those related to riboflavin metabolism, indicating metabolic complexity could also increase during endosymbiont evolution (17). Given that protists, exemplified by amoebae, represent primary hosts for chlamydiae in environmental settings, these findings underscore the critical role of amoebae in shaping the evolution of chlamydiae.

## Materials and Methods

### Sample Information

The wild amoeba clones used in this study are part of collection of over 700 amoebae isolated from soil or deer feces collected in various areas of the United States between 25-Sep-2000 and 13-Aug-2014 (Table S9). The 106 clones from this large collection were selected for this investigation in two parts. One set of 10 clones were selected for deep sequencing and investigation of the associated microbial community. To expand the analysis, another set of 96 clones were selected because they were previously reported to carry a Chlamydia symbiont in a PCR screen by Haselkorn *et al.* 2021 and because a whole genome sequence was already available (as part of an collaboration among the labs of authors J.B. Wolf, C.R.L. Thompson, D.C. Queller and J.E. Strassmann). Total DNA was extracted using the FastPrep-24 5G (MPbio, USA) and FastDNA Spin Kit for Soil (MPbio, USA). DNA concentrations were determined using a NanoDrop spectrophotometer (ND-2000, Thermo Scientific, USA) and stored at −20 °C for subsequent analysis.

### Whole Genome Sequencing and Processing

Genomic DNA was sequenced using Illumina paired-end reads. To maintain data integrity, we deposited the sequencing data in the National Center for Biotechnology Information (NCBI) Sequence Read Archive (SRA) database under accession number PRJNA975672 (Table S6). After removing host amoeba and food bacteria-related reads, we employed the remaining reads to detect and analyze associated chlamydiae (Figure S14).

We initially excluded reads from the host and food bacteria to recover high-quality metagenome-assembled genomes (MAGs) of amoeba symbionts. In brief, the raw data were first screened with FastQC (v0.11.5) to assess quality, followed by trimming and filtering of low-quality reads using Trimmomatic (v0.36, parameters: TruSeq3-PE.fa 2:30:10, LEADING 5, TRAILING 5, SLIDINGWINDOW 5:20, and MINLEN 30) (37). We aligned the trimmed reads to a single FASTA file containing the reference genomes of both *D. discoideum* AX4 (38) (GCF_000004695.1) and the food bacterium *Klebsiella pneumoniae* (GCF_000240185.1_ASM24018v2) (both downloaded from NCBI in June 2019) using bowtie2 (v2.4.5) (39). Mapped reads were used to identify the species of amoebae clones (Supplementary description 1). The 16s rDNA sequences from the clean data were extracted and assembled by SPAdes within phyloFlash (v3.4.1) (40), referencing to SILVA 138.1 (41). Only the 16S rDNA sequence larger than 1000 bp would be retained.

Unmapped reads were extracted, converted to fastq format with Samtools (v1.13) (42), and subjected to *de nov*o contig assembly with SPAdes (default parameters) within the metaWRAP pipeline (v1.3.2) (43). The contigs were binned, refined, and reassembled within the metaWRAP pipeline (43) using the binning module (parameters: –maxbin2 –metabat2), bin_refinement module (parameters: –c 20 –x 20), and the Reassemble_bins module (parameters: –c 50 –x 10), respectively. In addition to the two binners used within metaWRAP, we separately performed binning using Vamb (v3.0.8) (44). To ensure quality, we combined all MAGs and retained only MAGs with genome completeness exceeding 80% and contamination below 10%, as estimated by CheckM (v1.0.12) (45). The taxonomic classifications were assigned to MAGs via GTDB-Tk (Genome Database Taxonomy Toolkit (23), version=2.1.1, release R207).

### Population Clustering and Genomic Analyses

To assess the phylogenetic relationships of our chlamydiaceae MAGs among Chlamydiae, we downloaded reference genomes from GTDB (https://gtdb.ecogenomic.org/), collecting related information, including GC ratios, genomic sizes, completeness, contamination, and their hosts, if any (Supplementary description 2). We employed the tool dRep (v3.2.2) (46) to cluster draft MAGs into species-level clusters based on 95% average nucleotide identity (ANI) (parameters: -sa 0.95) (47). The genomic ANI of the reference genomes and our assembled MAGs was calculated using pyani (v0.3) (48). To explore the distribution of the chlamydiae that we found, the inStrain pipeline (v1.0.0) (49) was performed, and genomic completeness was used to represent their distribution (Supplementary description 3). Here, genomic completeness measures how much of a region is covered by sequencing reads and is calculated as the percentage of bases in a region that are covered by at least a single read. We only considered MAGs with genomic completeness greater than 50% and coverage > 1.

The open reading frames (ORFs) of each MAG were predicted using Prodigal (v2.6.3) (50). Functional annotation of Chlamydia was performed against the Kyoto Encyclopedia of Genes and Genomes (KEGG) database (51) using KofamScan (v1.3.0) (52). The membrane proteins were annotated against the Transporter Classification Database (TCDB) (53) and UniRef90 (54) by DIAMOND (v2.1.8.162) (55) (e-value < 10^-5^).

To construct a phylogenetic tree, we extracted and aligned the 120 conserved proteins from respective genomic set using GTDB-Tk (classify_wf, default parameters). the phylogenetic tree was constructed using IQ-TREE (v 2.2.2.7, parameters: -st AA -B 1000 -alrt 1000, -m MFP) (56) based on extracted and aligned conserved protein sequences and rooted using non-reversible model (57). Finally, visualized using Interactive Tree Of Life (iToL) (58). Genome visualization was carried out in Proksee (59), including horizontal gene transfers (HGT) identified using Alien Hunter (60). Pseudogenes were identified using Prokka (61) (--compliant) and Pseudofinder (v2.0) (62) (the “Annotate” module). The insertion sequences (ISs) within genomes were found using ISfinder (63). Genomes of cyanobacteria were downloaded from GTDB, including GCF_015207825, GCF_000317675, GCF_001693275, and GCF_002356215. The phylogenetic tree containing the outgroup was constructed and visualized using the same method as descripting above.

### Transmission Electron Microscopy

Based on the novel chlamydiae distribution among the first sets of 10 amoeba samples, we selected samples containing chlamydiae for Transmission Electron Microscope (TEM) analyses to visualize chlamydiae. Four samples, including QS4_A1, QS68_A2, QS74_A3, and QS1080_A10 (with QS4 belonging to *D. giganteum* and the rest to *D. discoideum*), were selected. The amoeba spores were fixed, dehydrated, and embedded in Spurr’s resin. After sectioning and staining with uranyl acetate and alkaline lead citrate, samples were observed on a Hitachi Model H-7650 TEM.

### Statistical analysis

All statistical tests and data analysis were performed in R (v4.2.2) (64) and Python (v3.7.10, https://www.python.org/). The Analysis of Variance (ANOVA) was conducted in R (*aov* function). Data visualization was achieved using ggplot2 (v3.4.0) and matplotlib (v3.7.0) (65) unless otherwise specified. The heatmap was generated using the pheatmap package in R. Specifically, the populations of MAGs were clustered by ANIm in dRep with “scale=NA,” and the functional heatmap was clustered using “euclidean” and normalized by “scale=’row’”. were clustered by “euclidean” and normalized by “scale=’row’”. The metabolic potential and graphical abstract were drawn based on our results (Supplementary description 4).

## Data availability

Sequencing data were deposited at the National Center for Biotechnology Information (NCBI) Sequence Read Archive (SRA) database with accession number PRJNA975672. The metagenome-assembled genomes of the three novel chlamydia were deposited at NCBI with BioSample IDs SAMN35628208 (*Ca. Dictychlamydia* sp. LF1), SAMN35621451 (Ca. *Dictychlamydia Ca. sp.* LF2), SAMN35628315 (*Ca. Feichlamydia* sp. LF3) and belong to BioProject ID PRJNA975672.

## Acknowledgments

We thank the members of our lab groups for their helpful comments. This material is based upon work supported by the Guangdong Natural Science Funds for Distinguished Young Scholar (2023B1515020096), the Innovation Group Project of Southern Marine Science and Engineering Guangdong Laboratory (Zhuhai) (311021006), the National Natural Science Foundation of China (31970384), and U.S. National Science Foundation grants DEB-1753743 and DEB-2237266.

