## Supplementary material for "Novel symbionts reveal amoebae as significant hosts for environmental chlamydiae": SI

**This file includes:**

Supplementary description

Supplementary Description 1-4

Supplementary Figure:

Figures S1 to S13

**Supplementary methods**

**Supplementary description 1**

We used a gene sequence to identify the species of each amoebae sample. This sequence should be detected in all samples and should be a conserved protein. Based on this, we extracted the mapped reads that mapped to the ribosomal RNA of the reference genome (GCF_000004695.1) (1), and assembled them by IDBA-UD (2). The contigs were blasted against the NCBI-nt database. Then, the assembled contigs were blasted against the NCBI-nt database (Nucleotide Sequence Database) for assigning the taxonomy.

**Supplementary description 2**

There are 768 genomes under *Chlamydiota* in GTDB, and 746 of them belong to *Chlamydiales* (order). All chlamydiae that the hosts were known belong to this order, including *Chlamydiaceae* (466 genomes, vertebrate-associated), *Rhabdochlamydiaceae* (68 genomes, arthropod-associated), *Parachlamydiaceae* (49 genomes, amoeba-associated), *Parachlamydiaceae_A* (2 genomes, amoeba-associated), *Simkaniaceae* (15 genomes, amoeba-associated), and *Waddliaceae* (5 genomes, amoeba-associated). The rests of them are hosts-unknown, that are called as environmental chlamydia. All of the 768 genomes were belonging to 160 species in GTDB with 95 % ANI cutoff, consistent with our MAG processing methods. We only selected the reference genomes for each population cluster based on providing in the database (release R207). This step generated 160 reference genomes from GTDB.

**Supplementary description 3**

As the description from inStrain (3) documentation: “A measure of how much of a region is covered by sequencing reads. Breadth is an important concept that is distinct from sequencing coverage and gives you an approximation of how well the reference sequence you’re using is represented by the reads. Calculated as the percentage of bases in a region that are covered by at least a single read. A breadth of 1 means that all bases in a region have at least one read covering them”. Here the breadth is the same as the genomic completeness for detecting the presence/absence of chlamydiae.

**Please note that this “genomic completeness” is not the same as “completeness” from CheckM (4).**

**Supplementary description 4**

The figures, that show the metabolic potential of the novel chlamydiae genomes, were drawn the draft picture in the Microsoft PowerPoint. The metabolic pathway were the collocted from the functional annonation against the KEGG datababse (5), and the potential transmembrane protein and exchange were supposed by the TCDB (6) and previous studies (7-11). The graphical abstract is drawn by hand according to the summary of the full text.

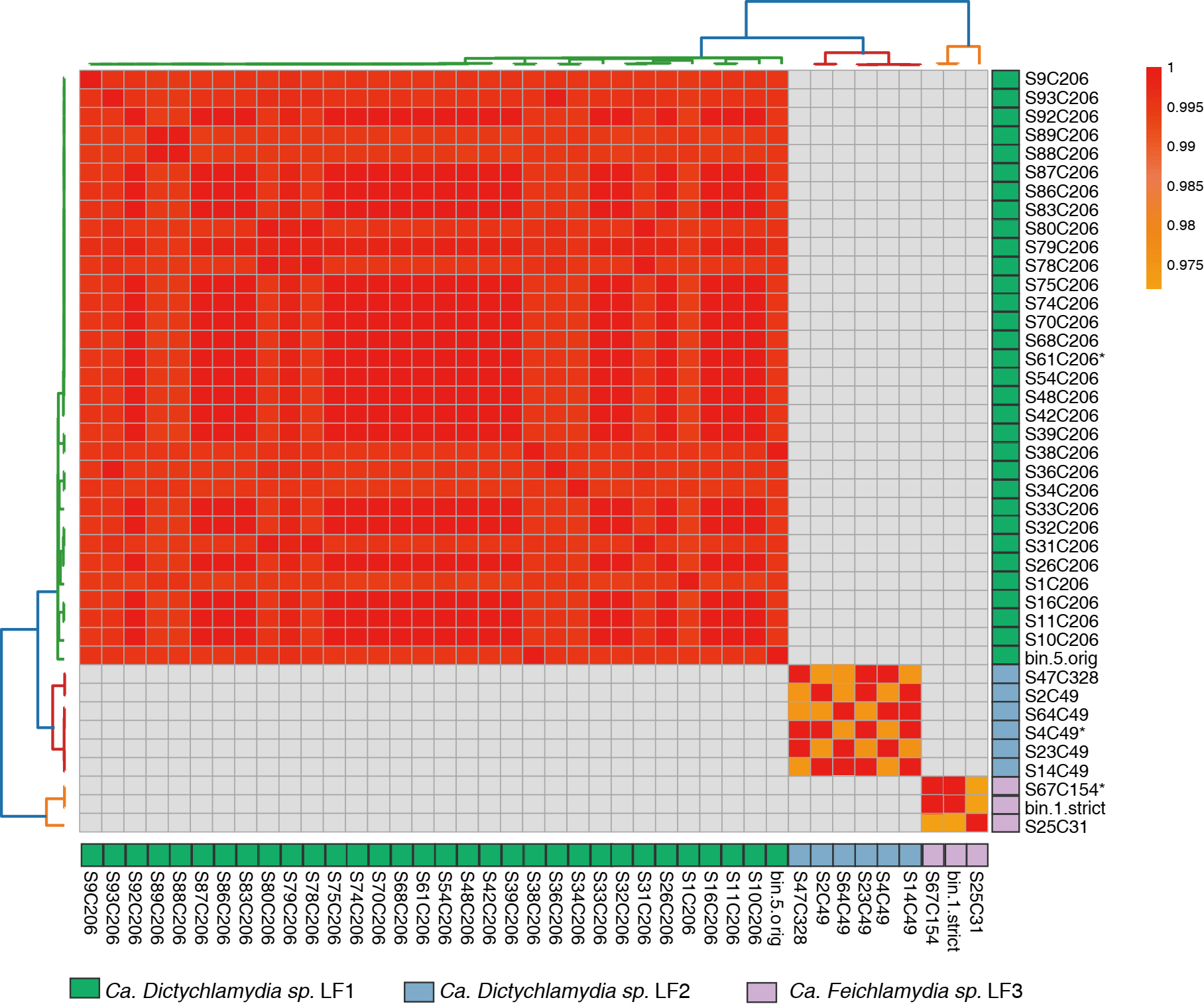

**Figure S1.** The population cluster of 41 chlamydiae genomes. At the 95% average nucleotide identity (ANI) cutoff and based on the 120 conserved proteins. The color blocks on the tree represent the species, including that the red is *Ca.* *Dictychlamydia sp.* LF1, the blue is *Ca.* *Dictychlamydia sp.* LF2, and yellow is *Ca.* *Feichlamydia sp.* LF3. The * is the genome that was selected for reference genomes based on the CheckM.

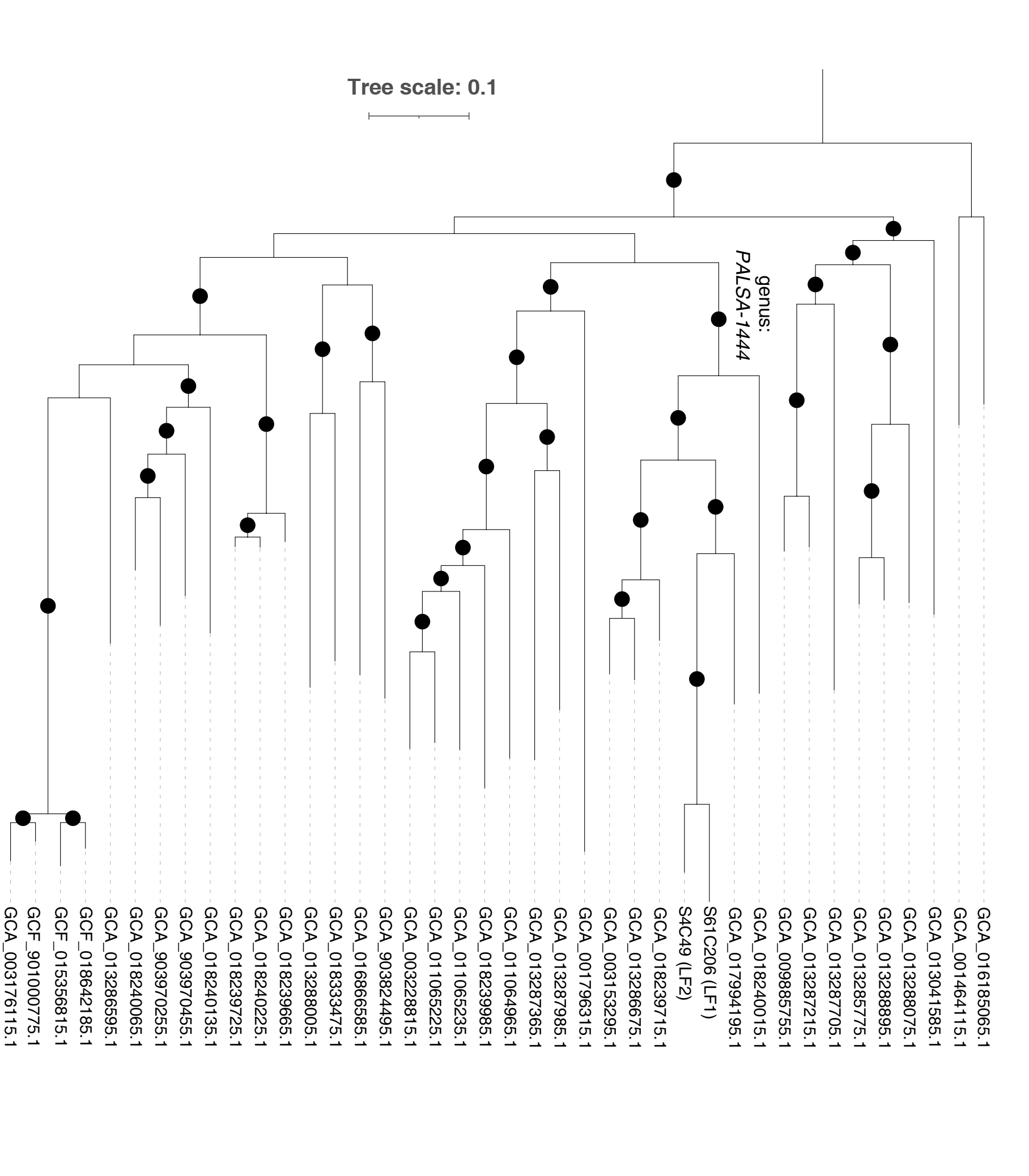

**Figure S2.** The phylogenetic tree shows relationships within *Rhabdochlamydiae*, containing 38 known species and the 2 *Ca. Dictychlamydia spp.* Identified in this study (*Ca. Dictychlamydia sp.* LF1 and *Ca. Dictychlamydia sp.* LF2) belonging to the environmental clade PALSA-1444. Substitution model: Q.insect+R6i. The black circles indicate branches with bootstrap values >70.

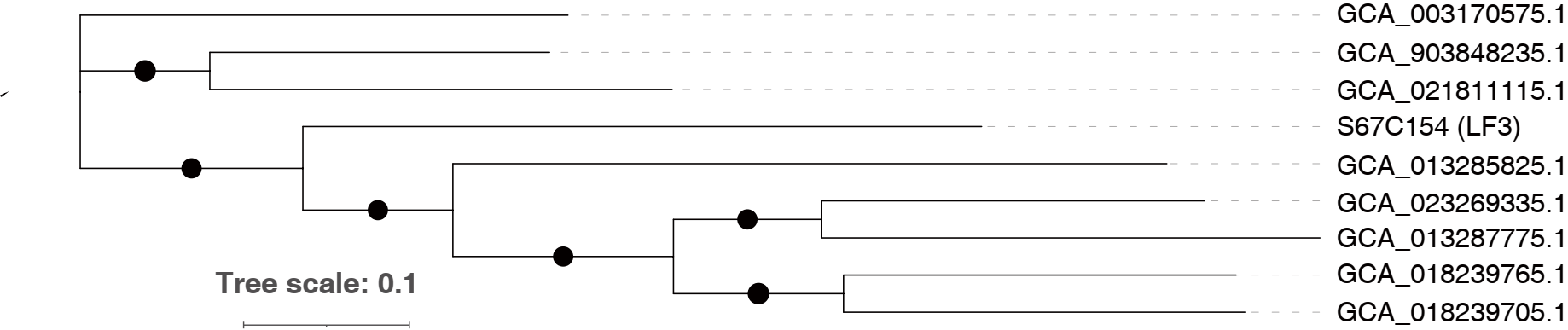

**Figure S3.** The phylogenetic tree within *FEN-1388*, containing 8 known environmental chlamydiae species and *Ca. Feichalmydia sp.* LF3 identified in this study. Substitution model: Q.insect+R6i. The black circles indicate branches with bootstrap values > 70.

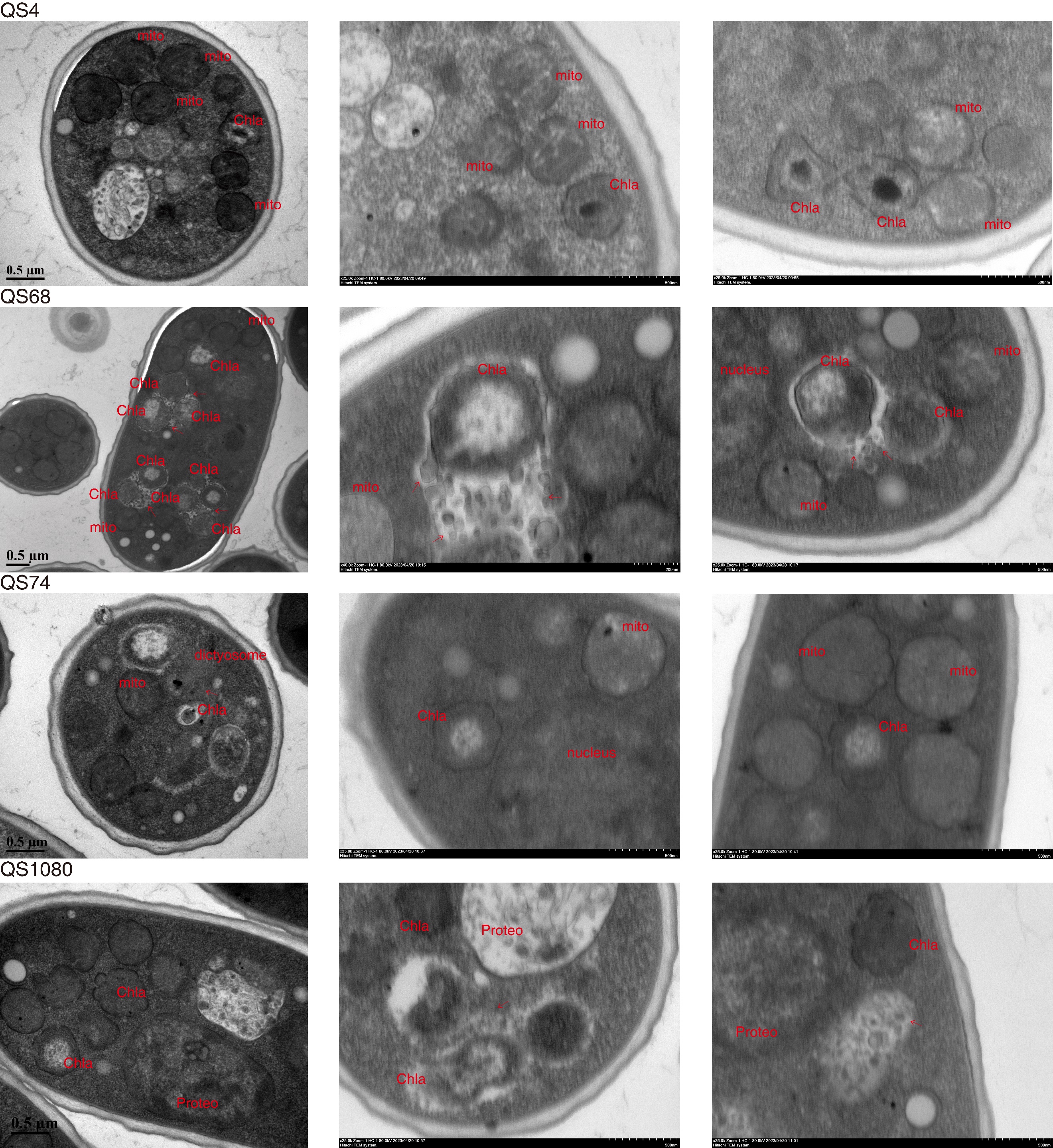

**Figure S4.** The chlamydiae in amoeba spores. Proteo: proteobacteria. Chla: chlamydiae. Mito: mitochondrion. Golgi: Golgi or dictyosome. The arrows are vesicles. The scale is shown in the picture. In each sample, the image on the left is an individual spore, followed by a close-up image on the right of endosymbionts.

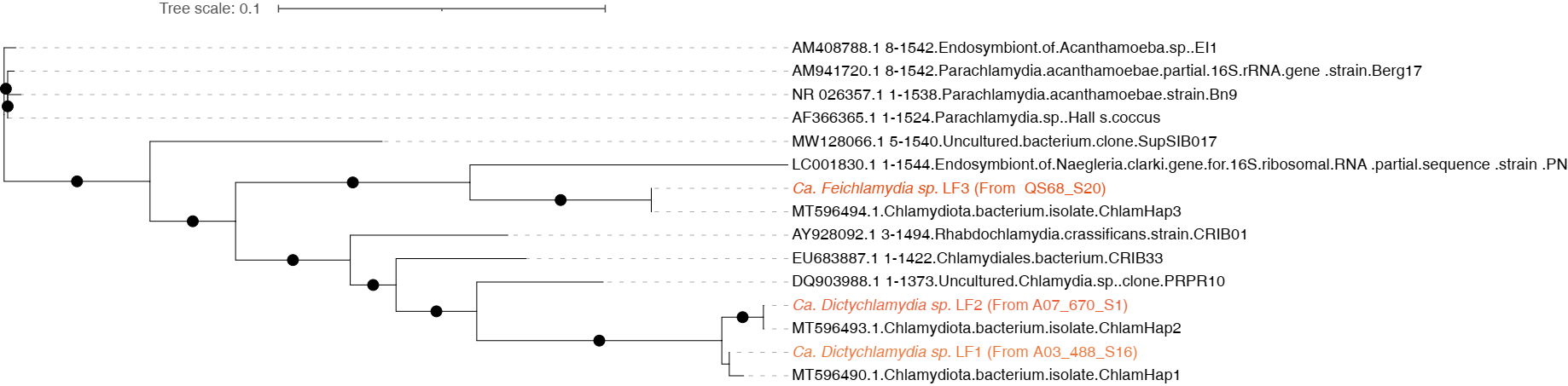

**Figure S5.** The tree plot shows the phylogenetic distance between the chlamydiae based on the 16S rDNA sequence. The black circles indicate branches with bootstrap values >70. Substitution model: Blosum62+F+G4.

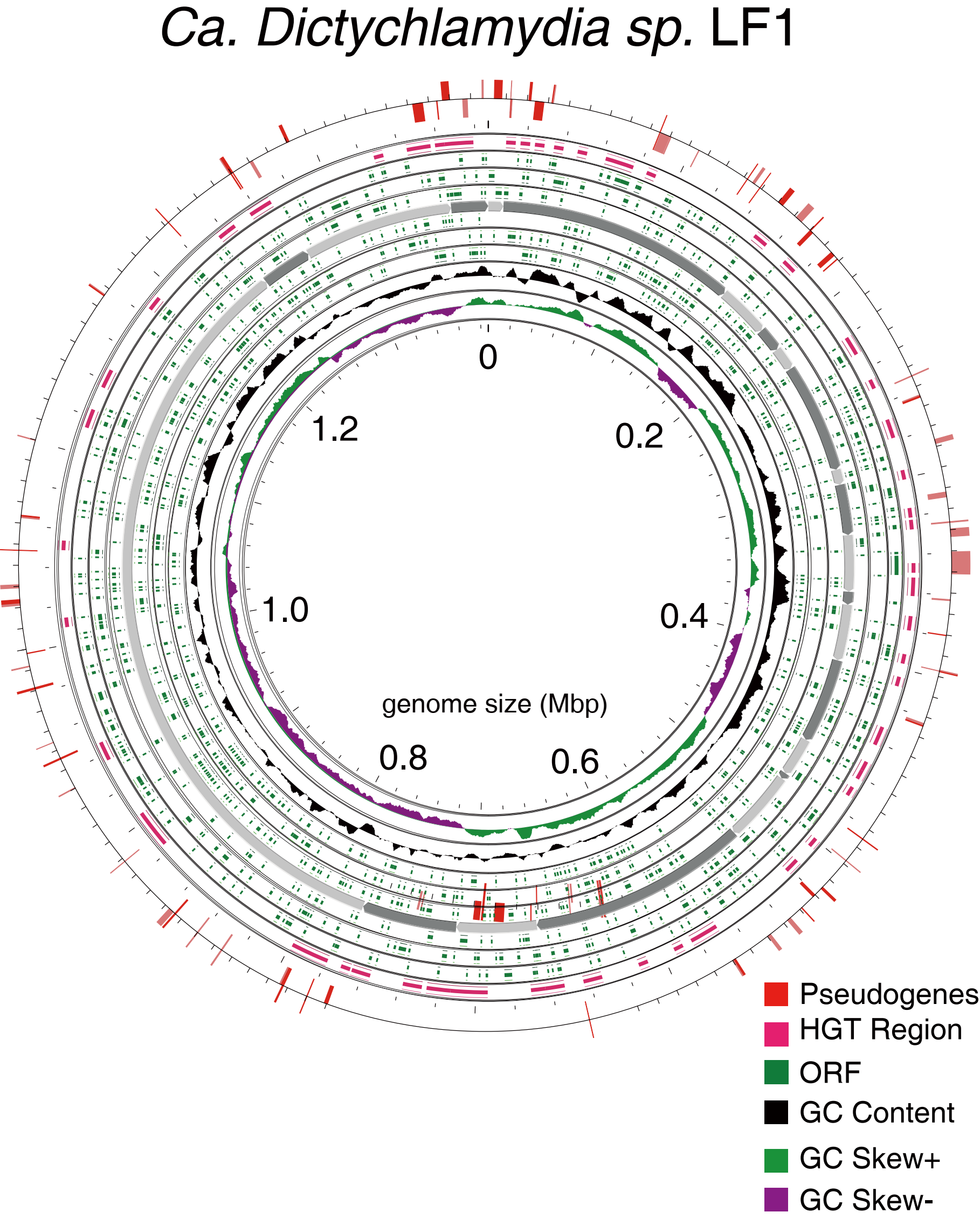

**Figure S6.** The genomic features of *Ca. Dictychlamydia sp.* LF1. From outside to inside, the circles are pseudogene regions (red), horizontal gene transfer (HGT) regions (pink), open reading frames (green), contigs (dark), GC content (black), and GC skew (green and purple).

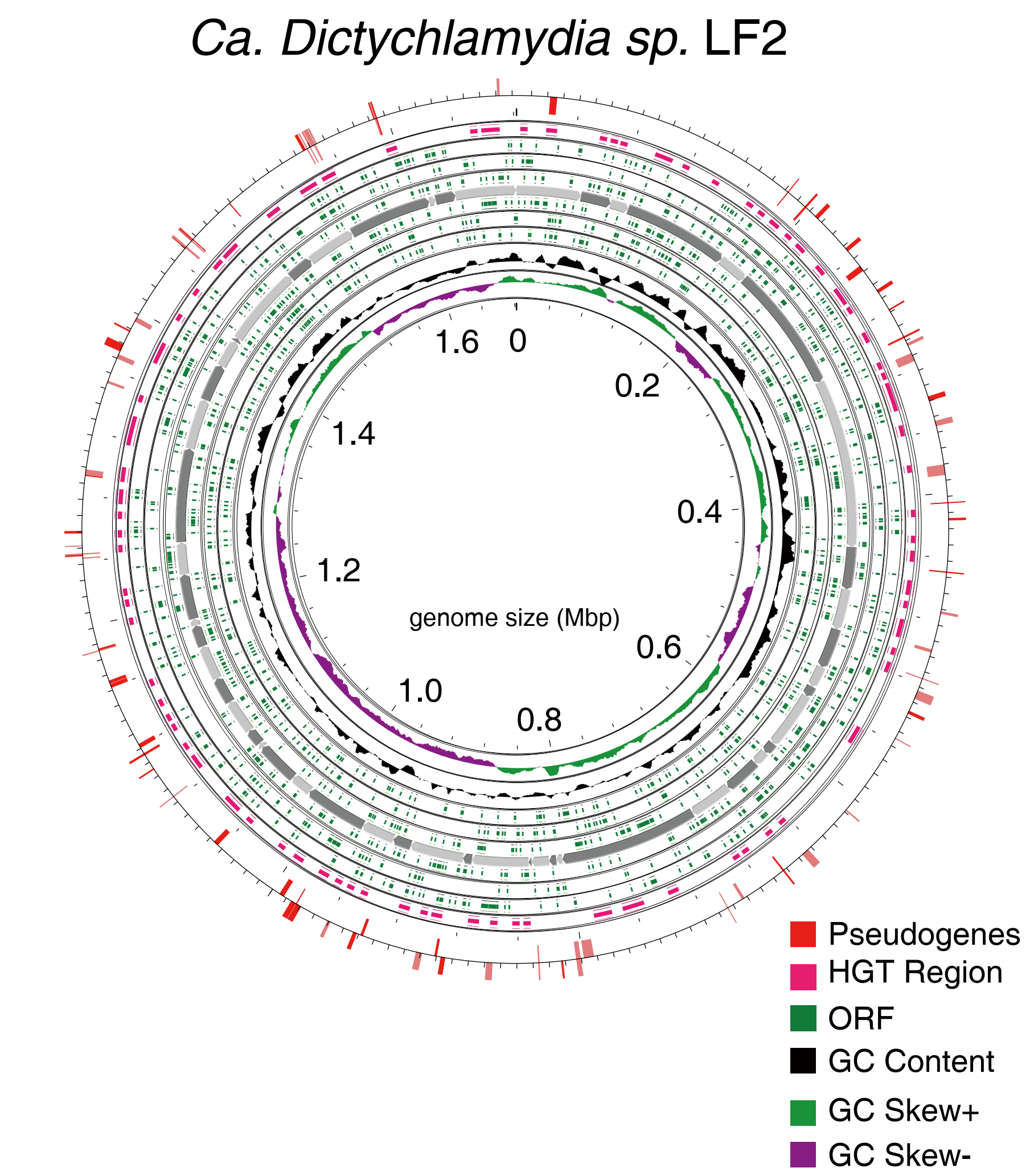

**Figure S7.** The genomic features of *Ca. Dictychlamydia sp.* LF2. From outside to inside, the circles are pseudogene regions (red), horizontal gene transfer (HGT) regions (pink), open reading frames (green), contigs (dark), GC content (black), and GC skew (green and purple).

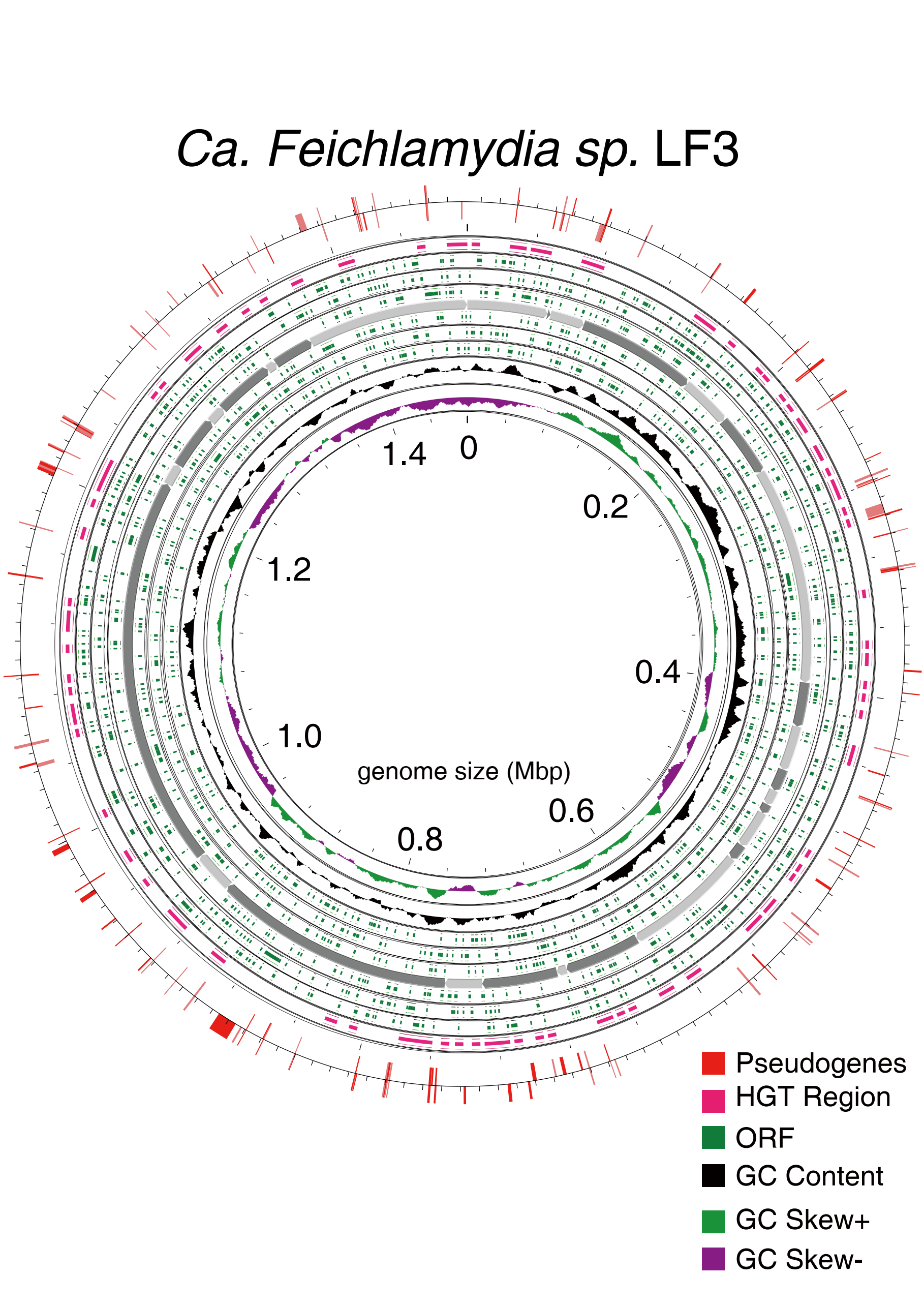

**Figure S8.** The genomic features of *Ca. Feichalmydia sp.* LF3. From outside to inside, the circles are pseudogene regions (red), horizontal gene transfer (HGT) regions (pink), open reading frames (green), contigs (dark), GC content (black), and GC skew (green and purple).

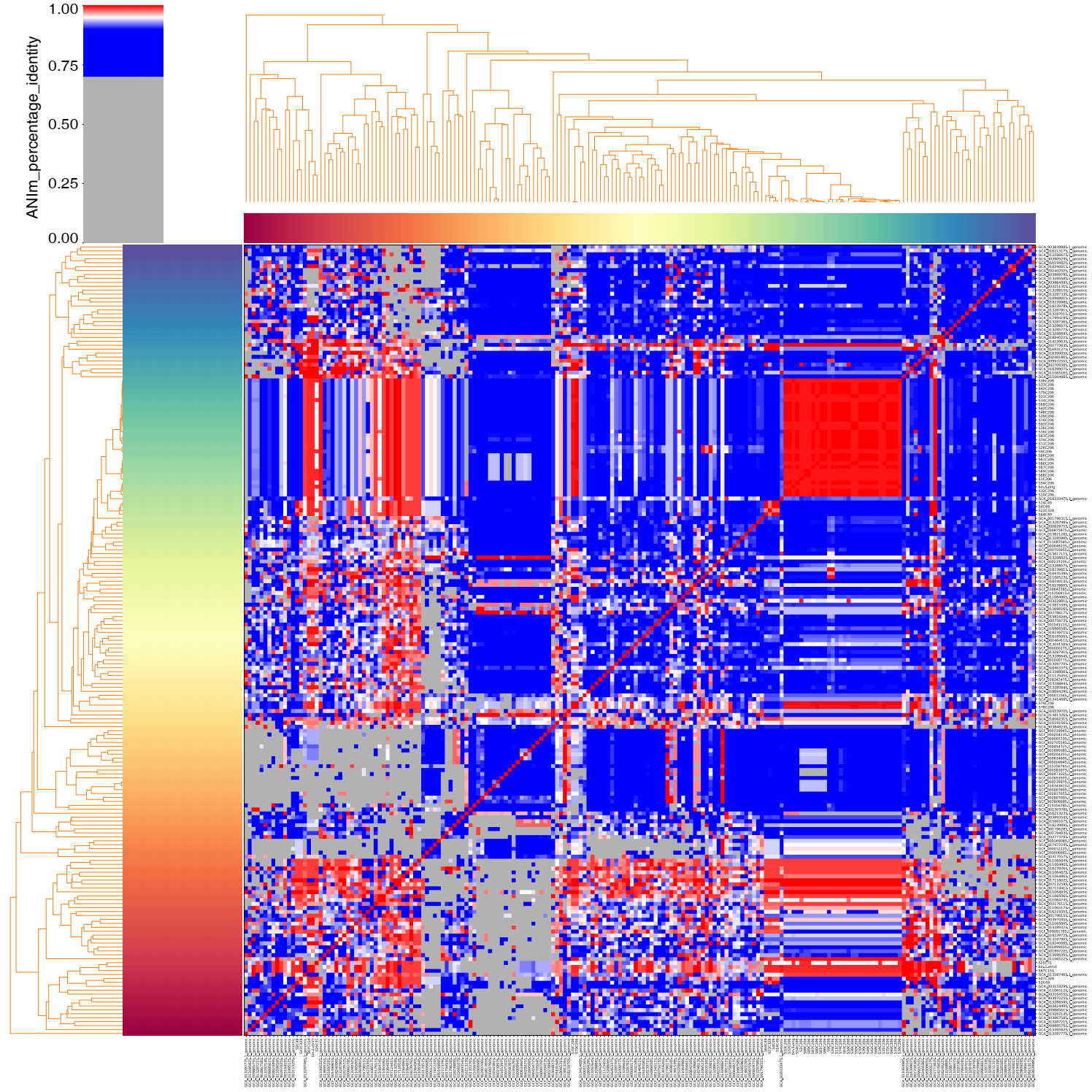

**Figure S9.** The genomic average nucleotide identity (ANI) across 160 chlamydia species and 41 novel chlamydia genomes. ANIs between 100% and 70% are shown from red to blue, and the rest of them are shown in gray.

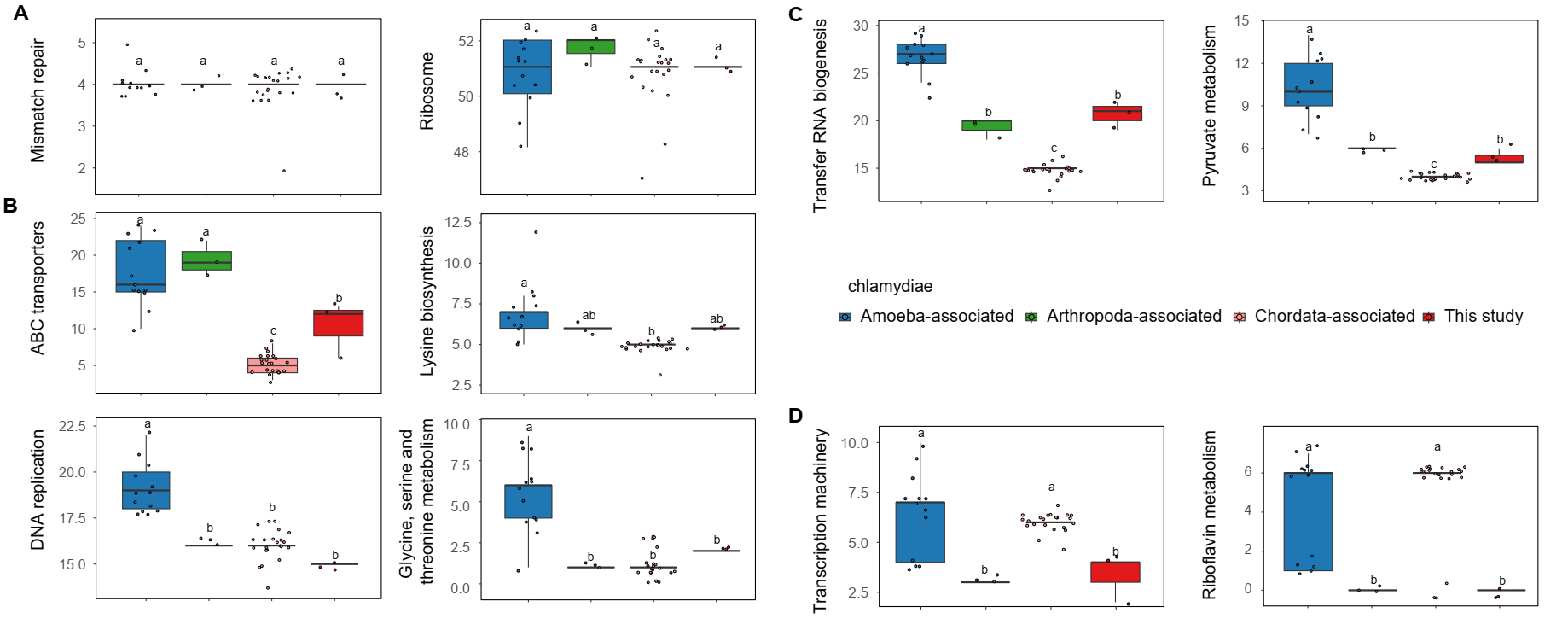

**Figure S10. A:** The boxplots show the parts of the functional categories of known-host chlamydiae were not shown significantly different between different hosts. **B:** The boxplots show the parts of the functional categories of known-host chlamydiae were shown significantly different between amoeba-associated and arthropod or vertebrate-associated chlamydiae. **C:** The boxplots show the parts of the functional categories of known-host chlamydiae were significantly decreased from amoeba-associated to arthropod-associated chlamydiae to vertebrate-associated chlamydiae. **D:** The boxplots show the parts of the functional categories of known-host chlamydiae appear to increase from amoeba-associated to vertebrate-associated chlamydiae. The letter marks their distinctiveness (ANOVA, *p* < 0.05).

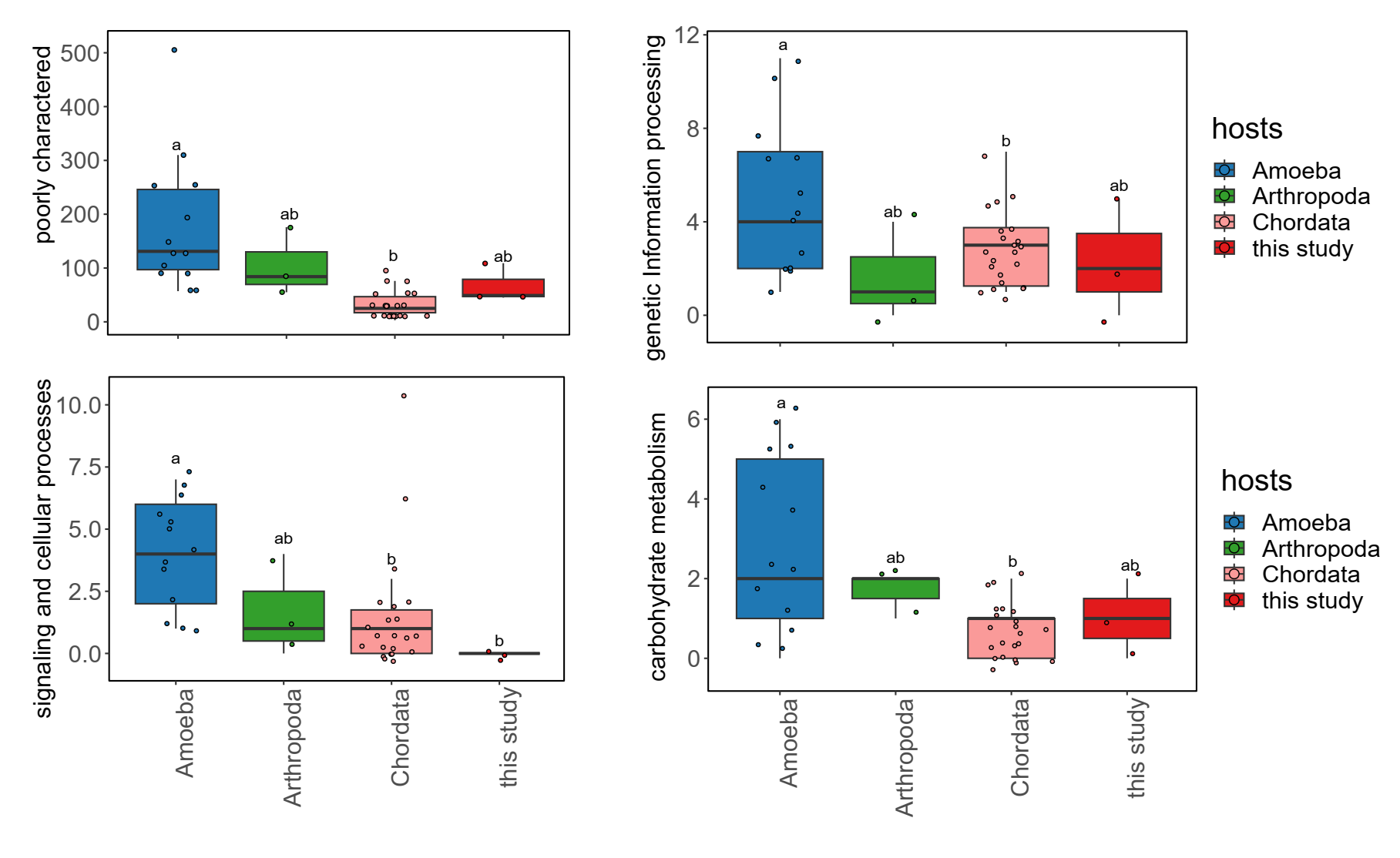

**Figure S11.** The boxplots show the parts of the functional categories of known-host chlamydiae were significantly decreased from amoeba-associated to arthropod or vertebrate-associated chlamydiae. The letter marks their distinctiveness (ANOVA, *p* < 0.05).

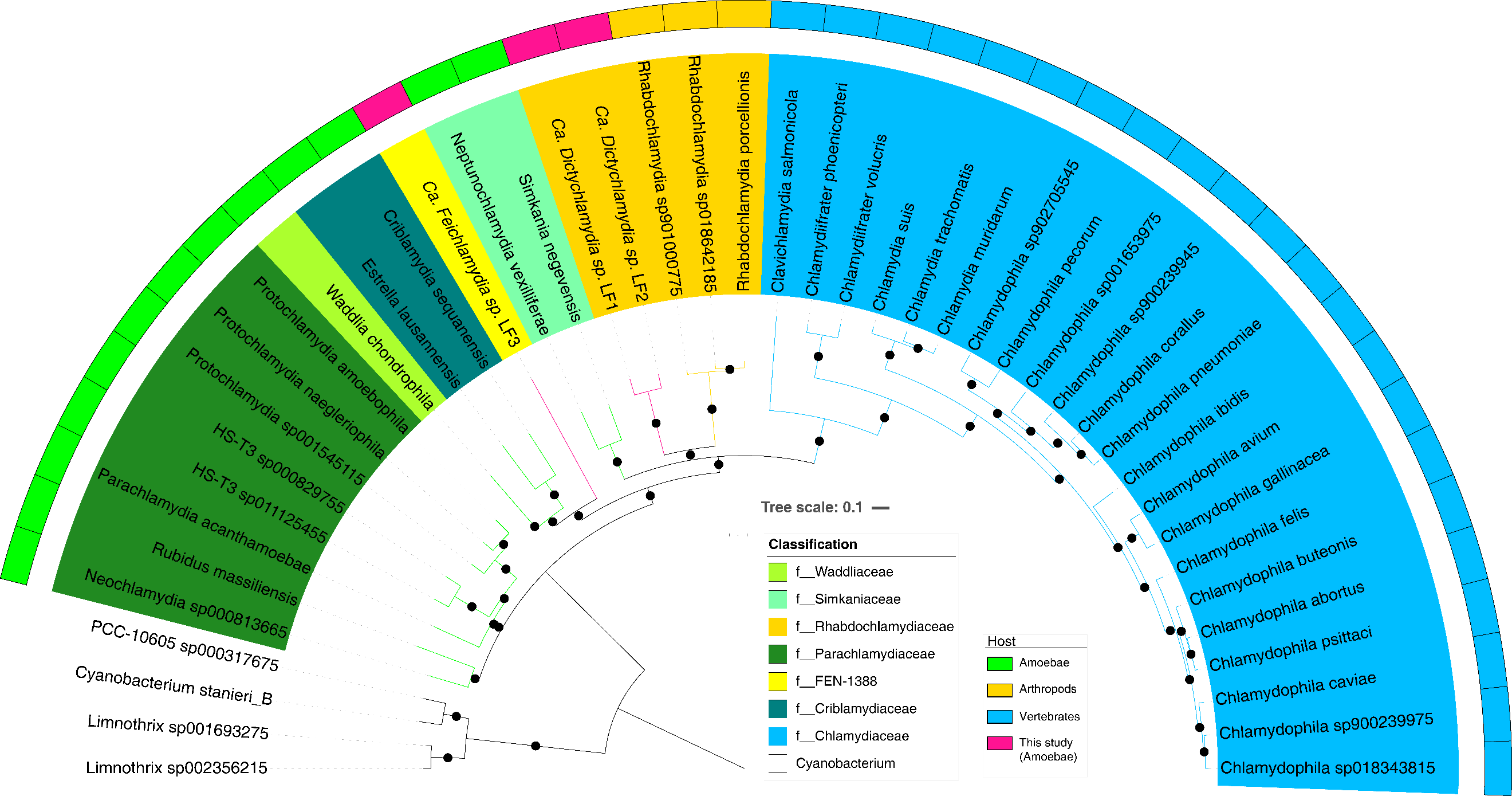

**Figure S12.** The phylogenetic tree of known hosting chlamydiae based on the 120 conserved proteins. Four cyanobacteria are the outgroup to root the tree. The color blocks represent their classification in the family level. The points are the bootstrap (>70) based on the IQ-tree result. The blocks outside tree represent the hosts of chlamydiae. Substitution model: Blosum62+F+R4.

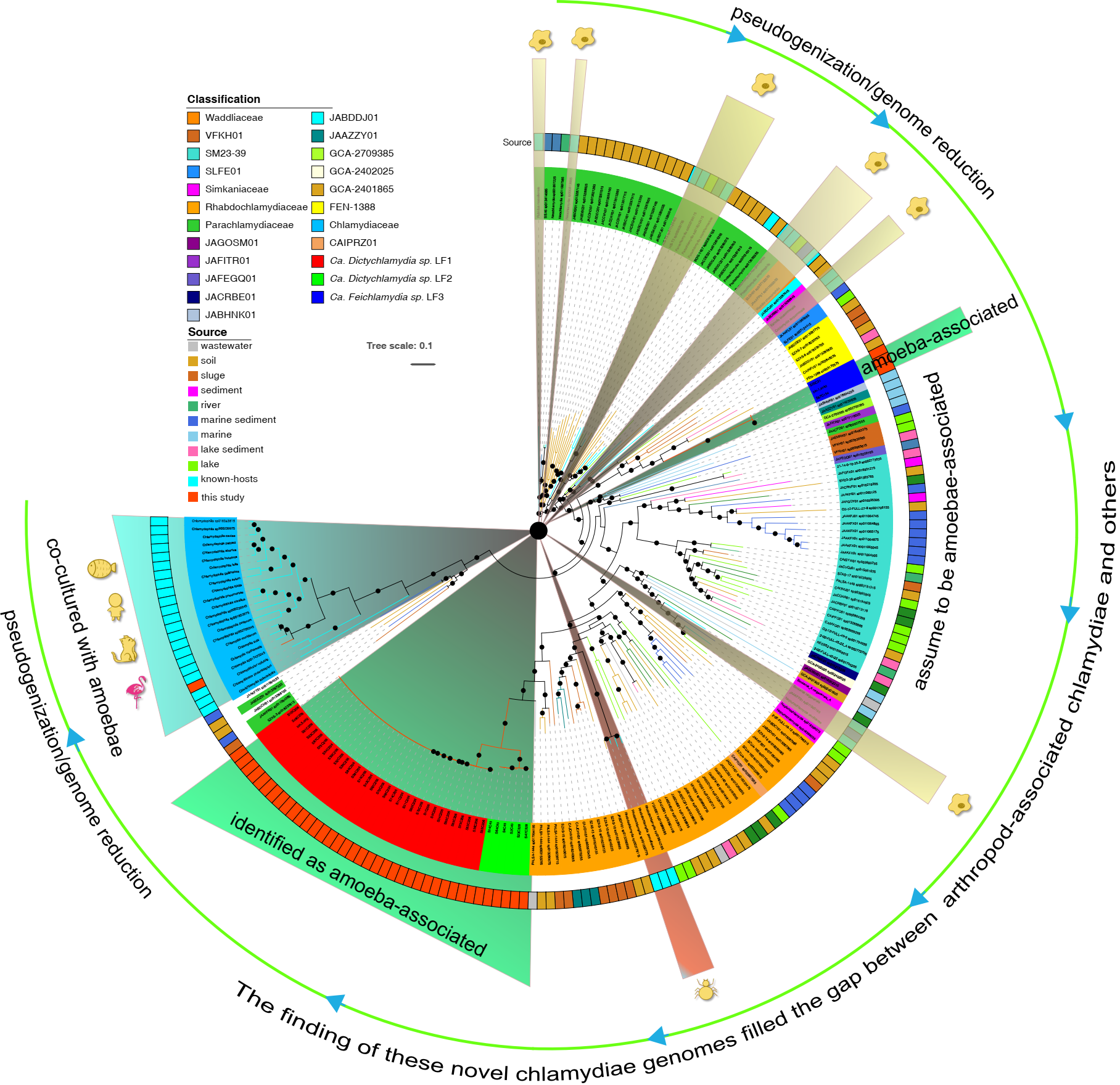

**Figure S13.** Inference of evolutionary processes based on the results and discussions. The phylogenetic tree is identical to Figure 1, which was rooted in amoebae-associated Chlamydiae. Following this transition, the characteristics of the genomes would be characterized by pseudogenization and genome reduction. Consequently, we bridged the gap between arthropod-associated Chlamydiae and other groups.

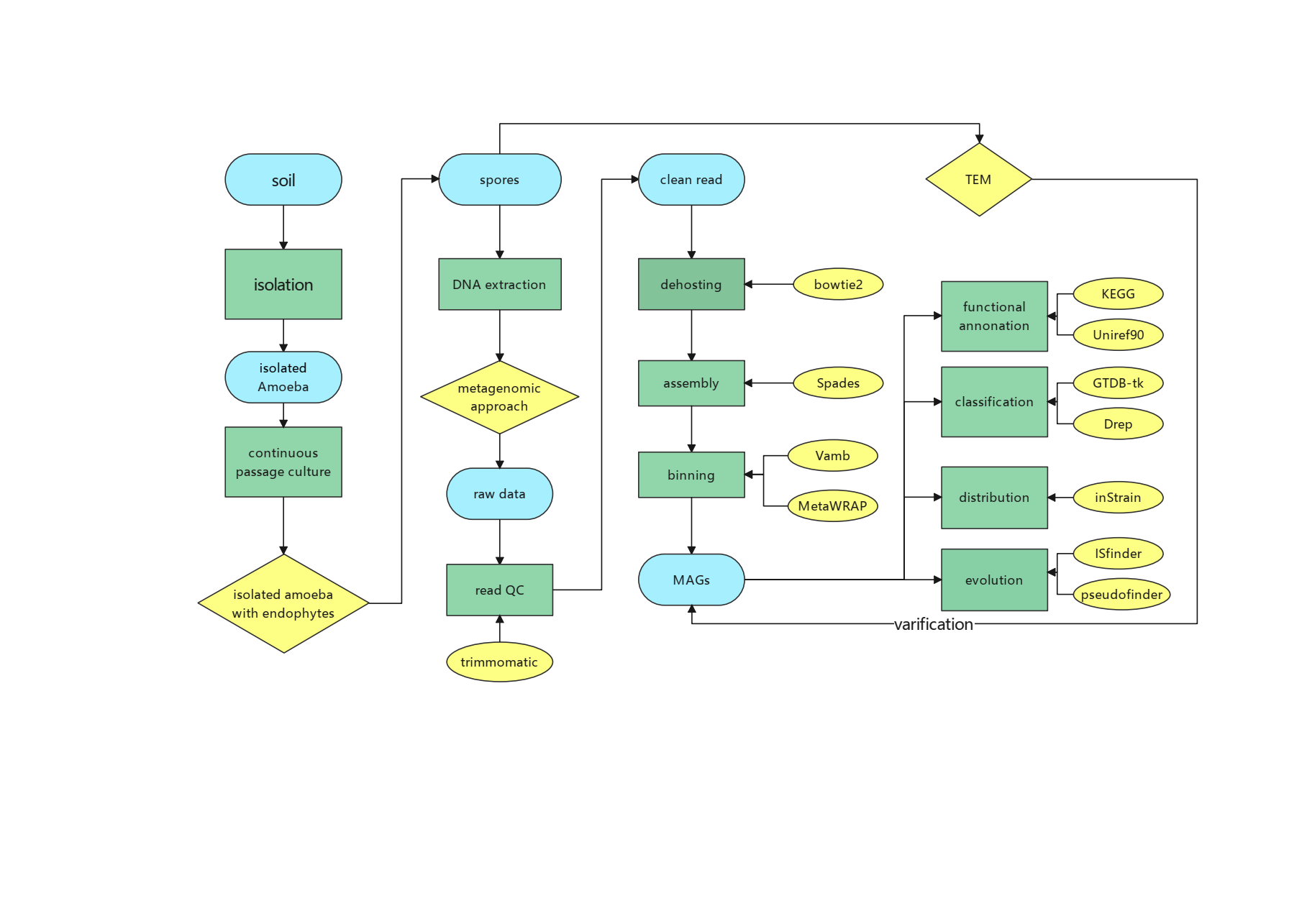

**Figure S14.**A brief workflow in this study. Stating from the sampling, we used multiple approaches for confirming our results.
